# Oncogenic BRAF Induces Whole-Genome Doubling Through Suppression of Cytokinesis

**DOI:** 10.1101/2021.04.08.439023

**Authors:** Revati Darp, Marc A. Vittoria, Neil J. Ganem, Craig J. Ceol

**Author notes:** Correspondence to: Craig J. Ceol, 368 Plantation Street, ASC-1041, Worcester, MA 01695.

## Abstract

Melanomas and other solid tumors commonly have increased ploidy, with near-tetraploid karyotypes being most frequently observed. Such karyotypes have been shown to arise through whole-genome doubling events that occur during early stages of tumor progression. The generation of tetraploid cells via whole-genome doubling is proposed to allow nascent tumor cells the ability to sample various pro-tumorigenic genomic configurations while avoiding the negative consequences that chromosomal gains or losses have in diploid cells. Whereas a high prevalence of whole-genome doubling events has been established, the means by which whole-genome doubling arises is unclear. Here, we find that BRAF^V600E^, the most common mutation in melanomas, can induce whole-genome doubling via cytokinesis failure *in vitro* and in a zebrafish melanoma model. Mechanistically, BRAF^V600E^ causes decreased activation and localization of RhoA, a critical cytokinesis regulator. BRAF^V600E^ activity during G1/S phases of the cell cycle is required to suppress cytokinesis. During G1/S, BRAF^V600E^ activity causes inappropriate centriole amplification, which is linked in part to inhibition of RhoA and suppression of cytokinesis. Together these data suggest that common abnormalities of melanomas linked to tumorigenesis – amplified centrosomes and whole-genome doubling events – can be induced by oncogenic BRAF and other mutations that increase RAS/MAPK pathway activity.

**Statement of Significance:** Whole-genome doubling is prevalent in many types of solid tumors and important in shaping tumor genomes, yet the causes of whole-genome doubling are not well understood. Here, we discover that oncogenic BRAF^V600E^ can induce whole-genome doubling through suppression of cytokinesis, and BRAF^V600E^-induced whole-genome doubling can occur in melanocytes and be present in nascent melanoma cells upon tumorigenesis.

## Introduction

Increased ploidy is a common feature of solid tumors. The most frequently observed increased karyotypes approach tetraploidy, which led to the hypothesis that such ‘near-tetraploid’ tumors had undergone a whole-genome doubling (WGD) event during tumor progression and subsequently experienced a small net loss of chromosomes (Carter et al., 2012; Dewhurst et al., 2014). Recent bioinformatic analyses support this hypothesis, showing that WGD events are prevalent in a diverse set of solid tumors, and nearly 37% of all solid tumors measured, including 40% of melanomas, experienced at least one WGD event in their progression (Quinton et al., 2021; Zack et al., 2013). Based on these analyses, WGD frequently occurs early in tumor formation, and the presence of tetraploid cells in some pre-cancerous lesions, such as Barrett’s esophagus and lesions of the cervix and kidney, suggests that WGD may even precede frank tumor formation in some tissues (Galipeau et al., 1996; Olaharski et al., 2006; Reid et al., 1996; Shackney et al., 1995) Tetraploidy was also observed in hyperplastic lesions of the pancreas (Tanaka et al., 1984), in localized prostate cancer (Deitch et al., 1993; Montgomery et al., 1990; Pihan et al., 2001) and some colon adenomas (Hamada et al., 1988; Levine et al., 1991), and for certain malignancies, such as oral tumors (Zaini et al., 2018), tetraploidy is a strong predictor of malignant transformation. Additionally, in established cancers from many tissue types WGD is a predictor of poor clinical outcome (Bielski et al., 2018).

Tetraploidy has been experimentally linked to tumorigenesis. Viral-induced cell fusion has been shown to enhance the transformation and tumor-forming capabilities of different cell types (Duelli and Lazebnik, 2007; Duelli et al., 2007; Gao and Zheng, 2010; Hu et al., 2009). Additionally, in mouse mammary epithelial cells that were made tetraploid through treatment with the actin filament poison dihydrocytochalasin B, tetraploid cells were able to form tumors in mice whereas their isogenic diploid counterparts were not (Fujiwara et al., 2005). In support of a role for WGD in tumorigenesis, deep sequencing of tumor samples has shown WGD to be an early event in non-small cell lung cancer, medulloblastoma and other tumor types (Carter et al., 2012; Dewhurst et al., 2014; Jamal-Hanjani et al., 2017; Jones et al., 2012).

There are different and mutually inclusive ways in which tetraploidy could contribute to tumorigenesis. First, tetraploidy can enable cells to become tolerant to the negative consequences of chromosome gains, losses, gene deletions, and inactivating mutations (Davoli et al., 2013; Hwang et al., 2021; Lopez et al., 2020; Passerini et al., 2016; Sheltzer et al., 2012; Thompson et al., 2006; Torres et al., 2008). Hence, tetraploidy is likely to allow tumor cells to withstand a higher incidence of mutations, thereby increasing the probability of adaptive changes. Second, tetraploid cells have an increased rate of chromosome missegregation (Mayer and Aguilera, 1990; Storchova et al., 2006; Wangsa et al., 2019), thus increasing the possibility that a developing tumorigenic clone will accumulate and tolerate the mutations needed for its progression to a malignant state (Davoli and de Lange, 2011). Thirdly, proliferating tetraploid cells are genetically unstable and can facilitate tumor progression by giving rise to aneuploidy, a known hallmark of cancer (Pfau and Amon, 2012).

Melanomas are a tumor type in which WGD is prevalent (Quinton et al., 2021). Although molecular genetic analyses have provided great insights into the genes that are involved in melanoma, very little is known about the process by which melanocytes with these lesions become tumorigenic, and whether any mutations underlie WGD in tumors is unclear. We examined melanocytes in zebrafish strains that are predisposed to melanoma and discovered an abundance of binucleate, tetraploid melanocytes. Tetraploidy was caused by expression of *BRAF^V600E^*, which increases RAS/MAPK-pathway activity and is commonly found in human melanomas. Using an *in vitro* model combined with live imaging, flow cytometry and immunofluorescence approaches, we found that BRAF^V600E^ generated tetraploid cells via cytokinesis failure and reduced activity of the small GTPase RhoA, which is critical for cytokinesis (Chircop, 2014). We also show that BRAF^V600E^ activity causes inappropriate centrosomal amplification, which is linked in part to the inhibition of RhoA and suppression of cytokinesis. Additionally, we show that zebrafish melanomas have a tetraploid karyotype and tumor-initiating cells in the zebrafish are tetraploid. These data collectively suggest that BRAF^V600E^-induced WGD occurs and has a role in tumor formation.

## Results

### *BRAF^V600E^* causes melanocytes in zebrafish to be tetraploid and binucleate

In this and previous studies, we used a zebrafish model of melanoma which combined melanocyte-lineage expression of human *BRAF^V600E^* with an inactivating mutation in the endogenous zebrafish *p53* gene (Ceol et al., 2011; Patton et al., 2005; Venkatesan et al., 2018). Animals of this genotype, *Tg(mitfa:BRAF^V600E^); p53(lf)*, develop melanomas that have histopathological and molecular features similar to those of human melanomas. To determine if tumors arising in this model exhibited ploidies consistent with having undergone a WGD event, we harvested tumors from *Tg(mitfa:BRAF^V600E^); p53(lf)* animals and quantified DNA content. The ploidy of these zebrafish melanomas was predominantly 4N and higher (Fig. 1A), indicating that WGD is likely a feature of this model. The analyzed tumors displayed a small fraction of 2N cells, which we speculate were admixed stromal cells.

**Figure 1:**
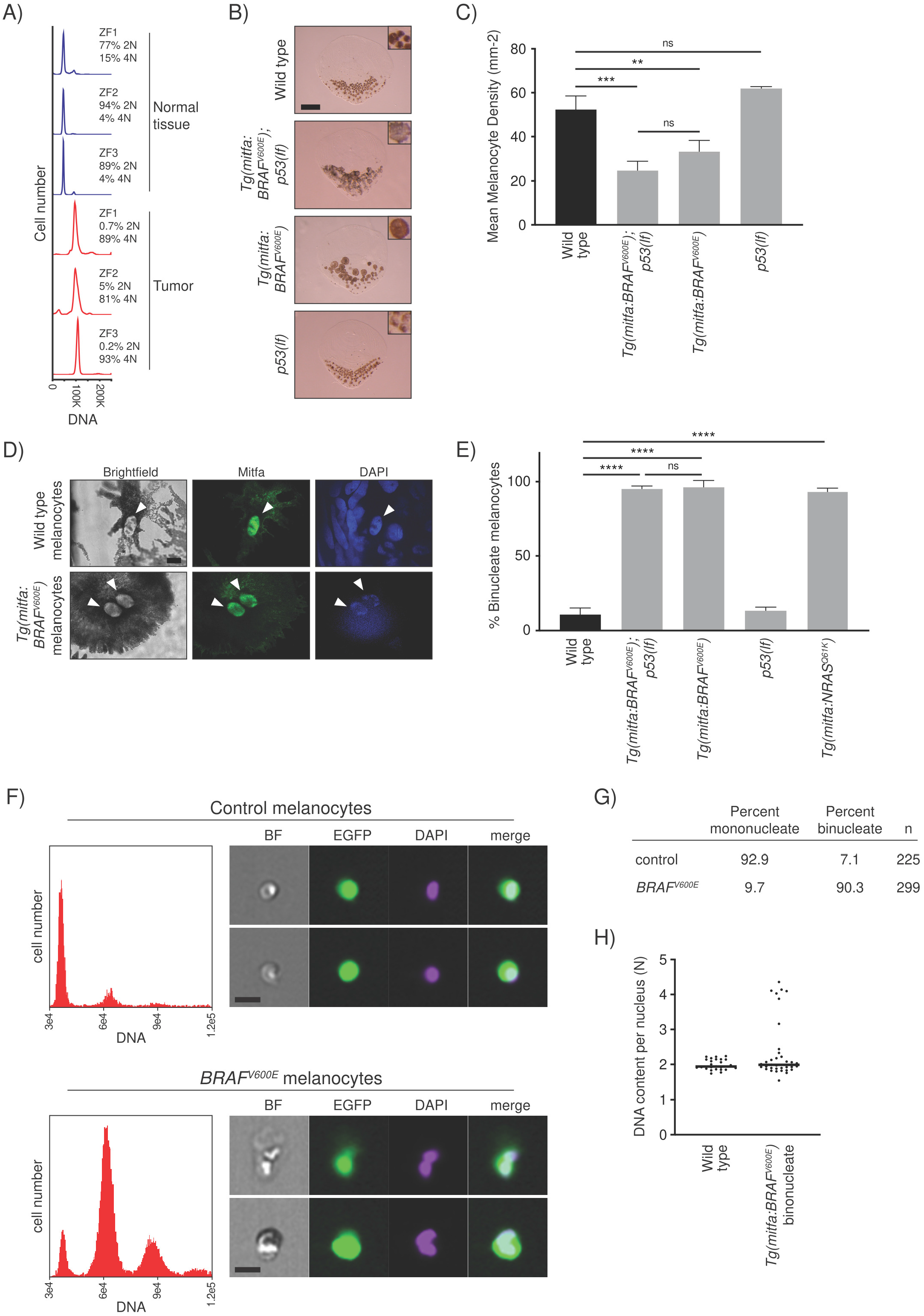
*BRAF^V600E^* causes melanocytes in zebrafish to be tetraploid. A) DNA content of normal and tumor tissue from *Tg(mitfa:EGFP); Tg(mitfa:BRAF^V600E^); p53(lf); alb(lf)* zebrafish. B) Scales from wild-type, *Tg(mitfa:BRAF^V600E^);p53(lf), Tg(mitfa:BRAF^V600E^) and p53(lf)* strains. Melanin pigment is dispersed throughout the cytoplasm of zebrafish melanocytes, revealing markedly different cell sizes. Scale bar = 250μm, insets are at same scale as one another. C) Quantification of melanocyte densities of wild-type, *Tg(mitfa:BRAF^V600E^);p53(lf), Tg(mitfa:BRAF^V600E^) and p53(lf)* strains. One-way ANOVA with Tukey’s multiple comparisons test, ***p<0.001, **p<0.01, ns = not significant. Error bars represent mean ± SEM. D) Images from brightfield (left), anti-Mitfa (middle) and DAPI (right) staining of a single wildtype (top) or *Tg(mitfa:BRAF^V600E^)* (bottom) epidermal melanocyte. Only the melanocyte nuclei stain positively for Mitfa. White arrowheads indicate nuclei within a single melanocyte. Scale bar = 5μm. E) Percent binucleate cells as determined by anti-Mitfa staining of pigmented melanocytes. One-way ANOVA with Tukey’s multiple comparisons test, ****p<0.0001, ns = not significant. Error bars represent mean ± SEM. F) Flow cytometry and DNA content analysis of control *Tg(mitfa:EGFP); alb(lf)* and *Tg(mitfa:EGFP); Tg(mitfa:BRAF^V600E^); alb(lf)* melanocytes with brightfield, EGFP and DAPI images of single melanocytes. G) Quantification of percent mononucleate and binucleate melanocytes from *Tg(mitfa:EGFP);alb(lf)* and *Tg(mitfa:EGFP); Tg(mitfa:BRAF^V600E^); alb(lf)* strains. Chi Square test, p=0.000009. H) DNA content analysis of wild-type and *Tg(mitfa:BRAF^V600E^)* melanocyte nuclei by confocal densitometry.

To investigate when the WGD event could occur, we began by examining melanocytes from *Tg(mitfa:BRAF^V600E^); p53(lf)* animals. We reasoned that closer analyses of epidermal melanocytes in the *Tg(mitfa:BRAF^V600E^)* strains might reveal the basis of the observed tetraploidy and provide insight into early cellular events that occur in melanoma tumorigenesis. To this end we developed assays to quantify and determine cell biological characteristics of these melanocytes. To quantify dorsal epidermal melanocytes, we treated fish with epinephrine then plucked and fixed scales to which these melanocytes are attached. As the number of melanocytes per scale depends on the size of the scale, we obtained a normalized melanocyte density measurement. Melanomas arise from dorsal regions of these zebrafish, and we found that the scale-associated epidermal melanocytes in these dorsal regions were larger in size and fewer in number than those of wild-type zebrafish (Fig. 1B, C). This was due to *BRAF^V600E^* expression, as *Tg(mitfa:BRAF^V600E^)* melanocytes were similar in size and number to *Tg(mitfa:BRAF^V600E^); p53(lf)* melanocytes, whereas *p53(lf)* melanocytes were similar to those of wild-type zebrafish. Previously, injection of *BRAF^V600E^* had been shown to cause nevus-like proliferations of melanocytes in zebrafish (Patton et al., 2005), and we also showed nevus-like melanocyte proliferations can arise in the *Tg(mitfa:BRAF^V600E^); p53(lf)* strain (Ceol et al., 2011). However, our current characterization of melanocytes in strains stably expressing the *Tg(mitfa:BRAF^V600E^)* transgene indicates that, aside from a few melanocytes that clonally proliferate, *BRAF^V600E^* expression primarily results in a reduced number and increased size of melanocytes. Cell size increases can be caused by increased ploidy, which could be reflected in a larger nuclear size (Sher et al., 2013). To determine if *BRAF^V600E^* expression caused nuclear enlargement, we stained for the melanocyte nuclear protein Mitfa. Most nuclei in large *Tg(mitfa:BRAF^V600E^)* melanocytes were similar in size to those of wild-type melanocytes; however, melanocytes in *Tg(mitfa:BRAF^V600E^)* contained two nuclei (Fig. 1D, E). Melanocytes from *Tg(mitfa:BRAF^V600E^); p53(lf)* animals were similarly binucleate, whereas *p53(lf)* melanocytes had one nucleus (Fig. 1E, Supplementary Fig. 1A), indicating that the binuclearity is associated with *BRAF^V600E^* expression.

To determine if the binuclearity we observed was uniquely associated with *BRAF^V600E^* or was caused by Ras/BRAF pathway overactivity in general, we stained scale-associated melanocytes expressing a common oncogenic variant of *NRAS*, mutations in which are present in about 28% of human melanomas (Cancer Genome Atlas, 2015). Melanocytes from animals expressing an *NRAS^Q61L^* oncogene that is commonly found in human melanomas were also binucleate (Fig. 1E, Supplementary Fig. 1B) (Dovey et al., 2009), indicating that binucleate cells arise from overactivation of RAS/MAPK signaling.

To examine ploidy of *Tg(mitfa:BRAF^V600E^)* melanocytes, flow cytometry and DNA densitometry were performed. Zebrafish melanocytes retain melanin pigment, so an *mitfa:EGFP* transgene and *albino* mutation were introduced so that melanocytes could be reliably identified by GFP-positivity and characterized without melanin spectral interference. Flow cytometry showed that *Tg(mitfa:BRAF^V600E^)* melanocytes were predominantly tetraploid, with small fractions of diploid and octoploid cells observed (Fig. 1F, G). DNA densitometry of fixed samples found that most nuclei in binucleate *Tg(mitfa:BRAF^V600E^)* melanocytes had a 2N DNA content (Fig. 1H). Therefore, expression of *BRAF^V600E^* causes melanocytes in zebrafish to become tetraploid as a result of having two nuclei, each with a 2N DNA content.

### BRAF^V600E^ binucleate, tetraploid cells arise via failure of cytokinesis

To determine how BRAF^V600E^ generates binucleate, tetraploid cells and to recapitulate the phenotype we observed in our zebrafish model, we developed an *in vitro* system suitable for mechanistic analyses. This system is based on a single-cell clone we created in which BRAF^V600E^ was inducibly expressed in RPE-1 FUCCI cells at a level similar to that of endogenous BRAF (Fig. 2A, B). We combined DNA content analysis with the fluorescent ubiquitin-based cell cycle indicator (FUCCI) reporter system to enable abnormal tetraploid cells in the G1 phase of the cell cycle (Cdt-mCherry-positive) to be distinguished from normal tetraploid cells in the G2 or M phases of the cell cycle (Geminin-GFP-positive) (Ganem et al., 2014; Sakaue-Sawano et al., 2008). In these cells, synchronization was performed by serum starvation, then after serum addition and progression through mitosis *BRAF^V600E^* was induced using doxycycline, and cells were synchronized again with a thymidine block. Following release from thymidine synchronization and progression through mitosis, G1 tetraploids were measured as Cdt-mCherry-positive cells with a 4N DNA content (Fig. 2C). Expression of *BRAF^V600E^* caused a nearly threefold increase in the percentage of G1 tetraploid cells (Fig. 2D). These cells were CyclinD1-positive, confirming that they were in G1 and not G2 cells that had dysregulated Cdt-mCherry expression (Supplementary Fig. 2A). Expression of wild-type or kinase-dead *BRAF* showed no similar increase in G1 tetraploids (Fig. 2D). Using a single-cell clone in which *BRAF^V600E^* was inducibly expressed in RPE-1 H2B-GFP cells, live-cell imaging was used to investigate whether the G1 tetraploids were binucleate and, if so, how they arose. Indeed, after *BRAF^V600E^* induction and release from synchronization, binucleate cells were observed in the following G1 phase of the cell cycle (Fig. 2E, F). These cells arose through failure of cytokinesis characterized by the formation then regression of the cytokinetic cleavage furrow. Modest increases in other mitotic defects, including lagging chromosomes, chromosome bridges and micronuclei, were observed although none of these increases was statistically significant (Supplementary Fig. 2B-D). Mitotic duration was not affected, even in cells that had undergone cytokinesis failure (Supplementary Fig. 2E). Together these data indicate that oncogenic BRAF^V600E^ causes WGD and the formation of binucleate, tetraploid cells by impairment of cytokinesis rather than cell fusion or other means.

**Figure 2:**
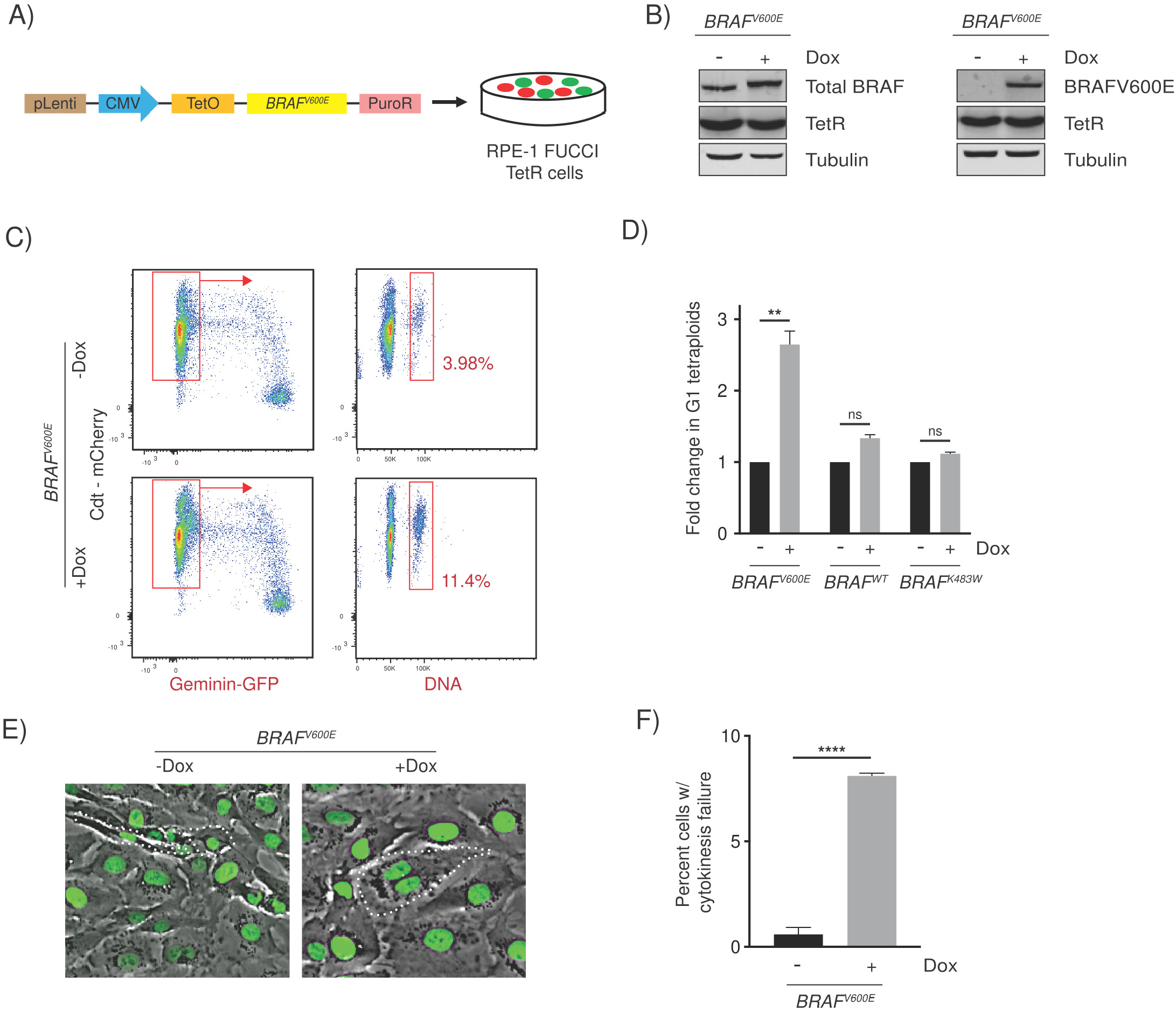
*BRAF^V600E^*-induced binucleate, tetraploid cells arise via cytokinesis failure. A) Generation of *BRAF^V600E^*-expressing RPE-1 FUCCI cell lines with a lentiviral-based doxycycline-inducible vector. B) Western blot showing inducible expression of BRAF^V600E^ (+Dox) using a BRAF and BRAF^V600E^-specific antibody. Expression of the Tet repressor protein is shown. Tubulin is used as the loading control. C) Flow cytometry plots of control (-Dox) and *BRAF^V600E^*-expressing (+Dox) cells. Tetraploid cells accumulating in G1 were quantified based on Cdt1-mCherry positivity and Hoechst incorporation. Percentages of G1 tetraploid cells in control and *BRAF^V600E^*-expressing cultures are indicated. D) Fold change in G1 tetraploids relative to the control (-Dox) are shown for *BRAF^V600E^, BRAF^WT^* and *BRAF^K483W^* (kinase-dead) -expressing cell lines. Fold change from 3 independent experiments is shown; unpaired Student’s *t* test, ** p < 0.01, ns = not significant. Error bars represent mean ± SEM. E) Merged phase contrast and GFP photomicrographs of H2B-GFP expressing control (-Dox) and *BRAF^V600E^*-expressing (+Dox) cells that have recently undergone mitosis. White dotted lines indicate 2 cells with 1 nucleus each that have separated following a successful cytokinesis (-Dox) and 1 cell with 2 nuclei that has failed cytokinesis (+Dox). F) Quantification of cytokinesis failure in H2B-GFP RPE-1 control (-Dox) and *BRAF^V600E^*-expressing (+Dox) cells. Percent cells with cytokinesis failure from 3 independent experiments is shown (total cells are n= 934 for -Dox and n=568 for +Dox). Unpaired Student’s *t* test, **** p< 0.0001. Error bars represent mean ± SEM.

### BRAF^V600E^ causes cytokinesis failure by reducing the localization and function of RhoA

To understand how BRAF^V600E^ inhibits cytokinesis, we investigated the localization and function of proteins that are central to the cytokinetic process. Proteins that have been extensively characterized as being critical during cytokinesis, mainly in contractile ring and cleavage furrow formation, are RhoA, a member of the RhoGTPase family and its scaffold protein Anillin (Chircop, 2014; Piekny and Glotzer, 2008). Following anaphase Anillin is required to maintain the assembly of cytokinetic furrow components at the equatorial cell cortex. In cells expressing *BRAF^V600E^*, Anillin staining was greatly reduced (Fig. 3A, B). Anillin localization is regulated by RhoA (Prokopenko et al., 1999), which activates and coordinates several downstream events in the cytokinetic process. RhoA is spatiotemporally activated and accumulates at the equatorial cell cortex in anaphase and during cytokinesis. This accumulation is both necessary and sufficient for cytokinesis to proceed (Basant and Glotzer, 2018). Similar to Anillin, RhoA localization to the cell equator was reduced in *BRAF^V600E^*-expressing cells (Fig. 3C, D). RhoA reduction in *BRAF^V600E^*-expressing cells was dependent on increased MAPK signaling because treatment with the MEK inhibitor trametinib or ERK inhibitor SCH772984 restored RhoA localization (Fig. 3C, D). Since RhoA localization is promoted by its activation at the equatorial cell cortex (Yuce et al., 2005), we quantified the levels of active, GTP-bound RhoA in *BRAF^V600E^*-expressing cells. Levels of GTP-bound RhoA were reduced as a consequence of *BRAF^V600E^* expression (Fig. 3E, F). If BRAF^V600E^ acts to reduce function of RhoA, then an increase in RhoA function would be predicted to suppress the effect of BRAF^V600E^ on the formation of binucleate, tetraploid cells. This occurred, as treatment of *BRAF^V600E^*-expressing cells with RhoA activators LPA and S1P reduced the formation of tetraploid cells (Fig. 3G). Furthermore, expression of the *RHOA^Q61L^* activated variant suppressed *BRAF^V600E^*-induced tetraploidy (Fig. 3H). These data indicate that BRAF^V600E^ reduces the activity of RhoA and its downstream effector Anillin, which underlies the failure of cytokinesis and the formation of binucleate, tetraploid cells.

**Figure 3:**
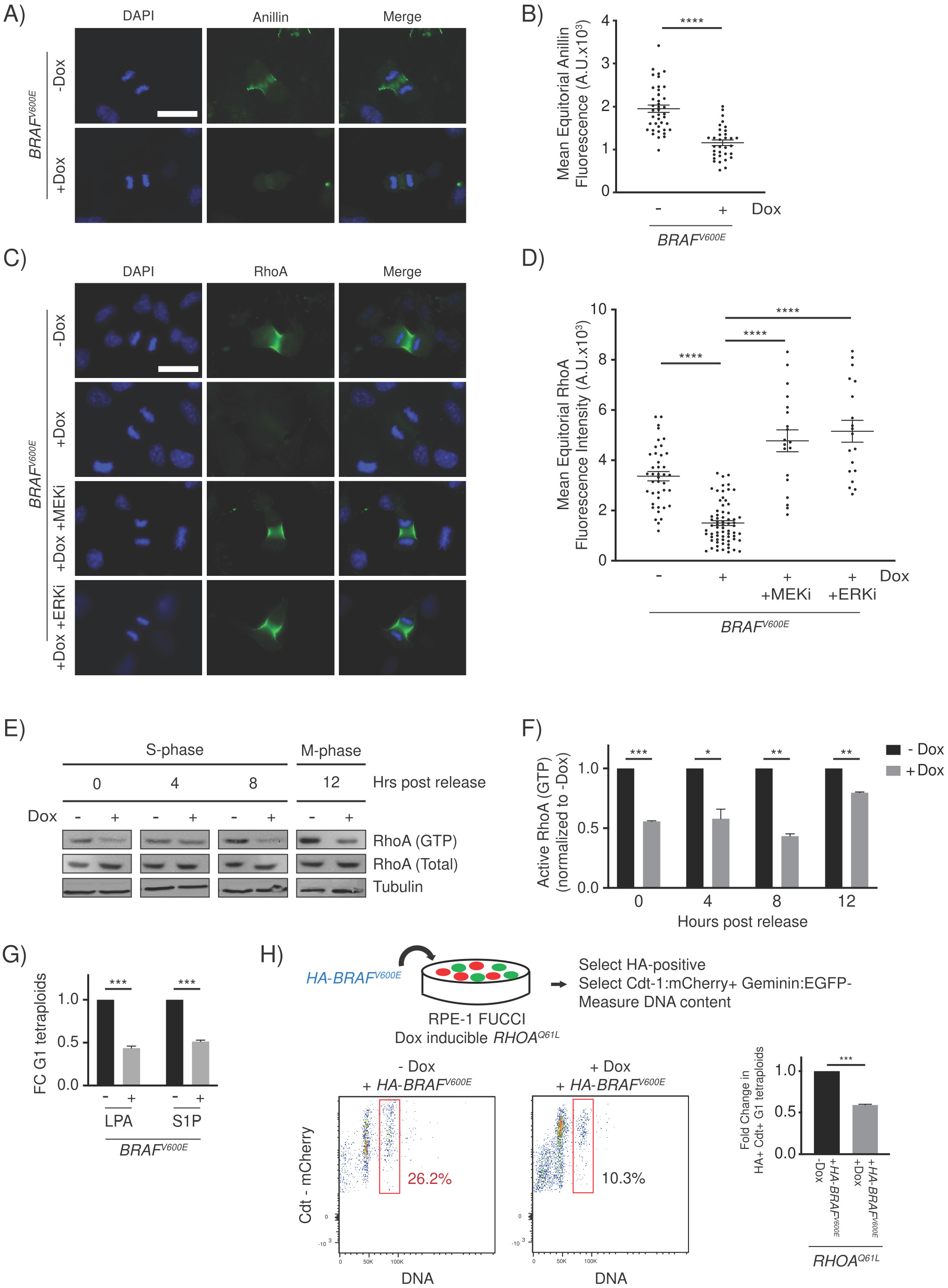
BRAF^V600E^ causes cytokinesis failure by reducing the localization and function of RhoA. A) DAPI and anti-Anillin staining in control (-Dox) and *BRAF^V600E^*-expressing (+Dox) anaphase cells. Images are maximum intensity projections of z-stacks. Scale bar = 7.5μM. B) Mean Anillin fluorescence intensity at the equator of control (n=80) and *BRAF^V600E^*-expressing (n=61) anaphase cells. Fluorescence intensities (mean gray values) of the equator were measured by sum intensity projections of z-stacks. Unpaired Student’s *t* test, ****p < 0.0001. Error bars represent mean ± SEM. C) DAPI and anti-RhoA staining in -*BRAF^V600E^* (-Dox) cells, *BRAF^V600E^*-expressing (+Dox) cells, and *BRAF^V600E^*-expressing (+Dox) cells treated with MEKi or ERKi. Drugs were added coincident with Dox administration. Images are maximum intensity projections of z-stacks (0.20μM). Scale bar = 7.5μM. D) Mean RhoA fluorescence intensity at the equator of -*BRAF^V600E^* (-Dox) (n=41), *BRAF^V600E^*-expressing (+Dox) cells (n=68), and *BRAF^V600E^*-expressing (+Dox) cells treated with MEKi or ERKi (n=21, n=21, respectively). Fluorescence intensities (mean gray values) of the equator were measured by sum intensity projections of z-stacks. One-way ANOVA with Tukey’s multiple comparisons test, **** p < 0.0001. Error bars represent mean ± SEM. E) Western blot analysis of immunoprecipitated RhoA-GTP from control (-Dox) and *BRAF^V600E^*-expressing (+Dox) RPE-1 cell lysates at different time points post thymidine release. Total RhoA protein and alpha tubulin were used as a controls. F) Western blot quantification of immunoprecipitated RhoA-GTP levels from (-Dox) and *BRAF^V600E^*-expressing (+Dox) RPE-1 cell lysates at different time points post thymidine release. Samples were normalized to the -Dox condition. Measurements from 3 independent experiments are shown. Unpaired Student’s *t* test, * p< 0.05, ** p < 0.01, *** p < 0.001. Error bars represent mean ± SEM. G) Fold change in G1 tetraploid cells following addition of RhoA activators. LPA (1μM) and S1P (1μM) were added coincident with DOX administration. Fold change in G1 tetraploids relative to the control (+Dox no drug) are shown. Measurements from 3 independent experiments are shown. Unpaired Student’s *t* test, *** p<0.001. Error bars represent mean ± SEM. H) G1 tetraploid generation following expression of HA-tagged-BRAF^V600E^ in *RHOA^Q61L^*-inducible cells. Experimental design (top): RPE-1 FUCCI cells with Dox inducible *RHOA^Q61L^* were transiently transfected with an HA-tagged-BRAF^V600E^-expressing construct and selected accordingly. G1 tetraploids were quantified (left) by gating HA-positive, Cdt-1:mCherry-positive cells with increased Hoechst incorporation. Fold change in G1 tetraploids (right), normalized to control (-Dox) cells. Unpaired Student’s *t* test, *** p < 0.001. Error bars represent mean ± SEM.

### BRAF^V600E^ and MAPK pathway activity is required during late G1 and early S phases for generating tetraploids

BRAF^V600E^ acts in G1/S to promote cell cycle progression, yet some reports have suggested that MAPK activity is important during mitosis (Liu et al., 2004; Mulner-Lorillon et al., 2017; Wright et al., 1999). To determine when BRAF^V600E^ and MAPK signaling is required to generate tetraploids, we treated *BRAF^V600E^*-expressing cells with the BRAF^V600E^ inhibitor vemurafenib, MEK inhibitor trametinib and ERK inhibitor SCH772984 at various points in the cell cycle and measured whether the inhibitors suppressed tetraploid formation (Fig. 4A). The ability of inhibitors to reduce downstream MAPK activity was confirmed by western blot of phosphorylated ERK (Supplementary Fig. 3A, B). Additionally, we also confirmed that treatment with inhibitors at the concentrations used did not cause cell cycle arrest and did not substantially impact growth kinetics, cell cycle progression or cell viability (Supplementary Fig. 3C-E). Treatment with inhibitors throughout the cell cycle suppressed formation of *BRAF^V600E^*-induced tetraploids (Fig. 4A). By contrast, treatment during the S/G2/M phases had little effect on tetraploid formation. Treatment during G1 and specifically during late G1 and early S suppressed formation of tetraploids. This suppression occurred with each of the three inhibitors tested as well as the ‘paradox-breaking’ BRAF^V600E^ inhibitors PLX7904 and PLX8394 (Supplementary Fig. 3F), which inhibit BRAF^V600E^ while not simultaneously activating downstream MAPK activity like vemurafenib does (Hatzivassiliou et al., 2010; Poulikakos et al., 2010; Zhang et al., 2015). Thus, BRAF^V600E^, and MAPK signaling in general, act during late G1 and early S phases to ultimately cause inhibition of RhoA and failure of cytokinesis.

**Figure 4:**
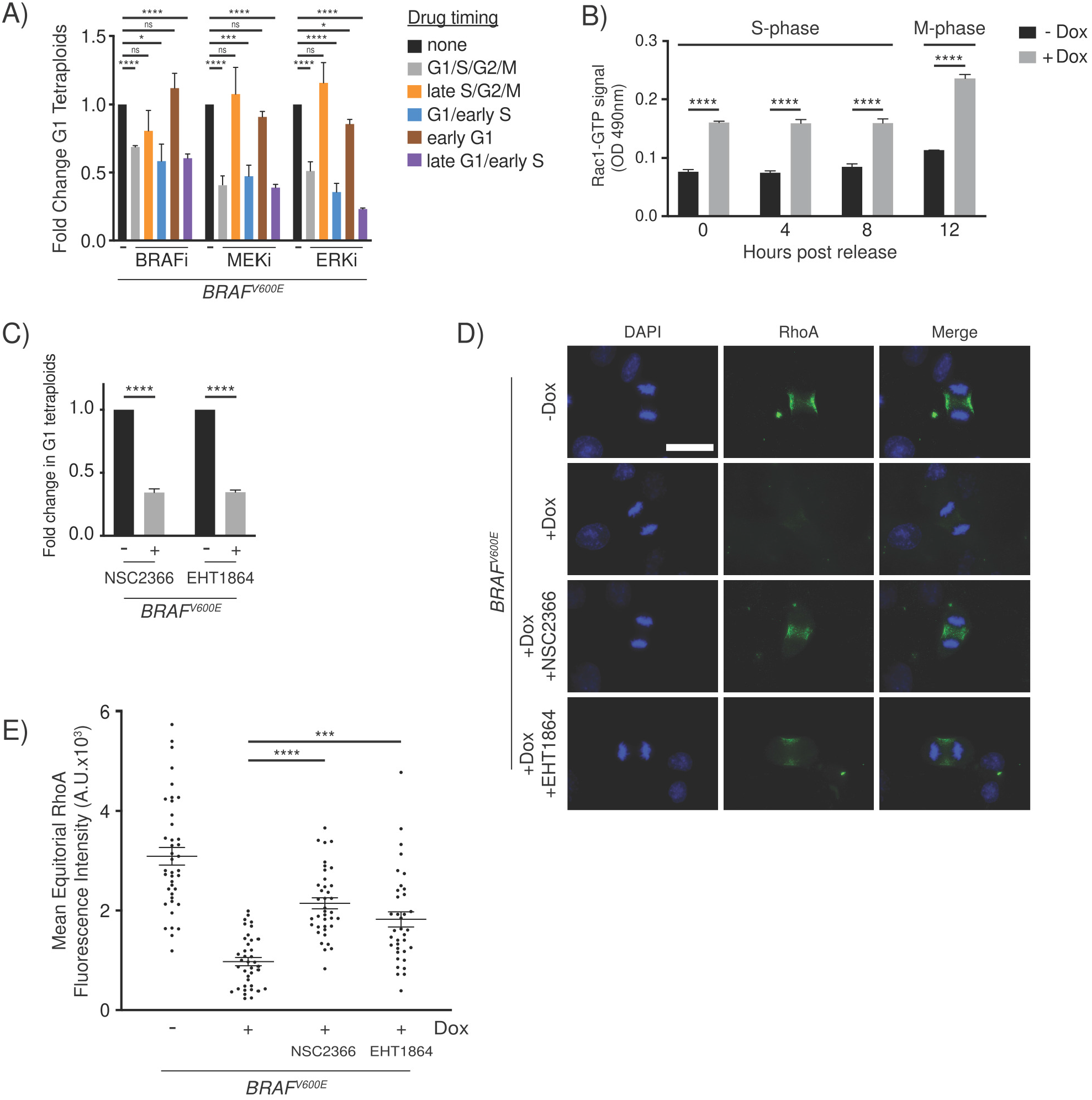
BRAF^V600E^ acts during G1/S and activates RAC1 to downregulate RhoA and generate tetraploid cells. A) Fold change in G1 tetraploids following inhibitor treatment. Fold changes are expressed relative to control (+*BRAF^V600E^*, no drug) cells. Fold change from 3 independent experiments is shown; unpaired Student’s *t* test, * p < 0.05, ** p < 0.01, ***p < 0.001, ****p < 0.0001, ns = not significant. Error bars represent mean ± SEM. B) ELISA-based quantification of RAC1-GTP levels in control (-Dox) and *BRAF^V600E^*-expressing (+Dox) RPE-1 cells. Cells were measured at the indicated timepoints post thymidine release. RAC-1 GTP signal was measured using a colorimetric assay at 490nM absorbance. Unpaired Student’s *t* test, **** p < 0.0001. Error bars represent mean ± SEM. C) Fold change in G1 tetraploids following addition of RAC1 inhibitors. NSC2366 and EHT1864 were added coincident with *BRAF^V600E^* induction. Fold changes are expressed relative to control (+*BRAF^V600E^*, no drug) cells. Fold change from 3 independent experiments is shown; unpaired Student’s *t* test, **** p<0.0001. Error bars represent mean ± SEM. D) DAPI and anti-RhoA staining in -*BRAF^V600E^* (-Dox) cells, *BRAF^V600E^*-expressing (+Dox) cells, and *BRAF^V600E^*-expressing (+Dox) cells treated with NSC2366 or EHT1864. Drugs were added coincident with Dox administration. Images are maximum intensity projections of z-stacks (0.20μM). Scale bar = 7.5μM. E) Mean RhoA fluorescence intensity at the equator of -*BRAF^V600E^* (-Dox) (n=40), *BRAF^V600E^*-expressing (+Dox) cells (n=38), and *BRAF^V600E^*-expressing (+Dox) cells treated with NSC2366 or EHT1864 (n=38, n=36, respectively). Fluorescence intensities (mean gray values) of the equator were measured by sum intensity projections of z-stacks. One-way ANOVA with Tukey’s multiple comparisons test, **** p < 0.0001. Error bars represent mean ± SEM.

### Rac1 is activated by BRAF^V600E^ and contributes to RhoA downregulation

To understand how BRAF^V600E^ and MAPK signaling activity during G1/S could lead to downregulation of RhoA and failed cytokinesis, we considered regulators of cytokinesis that are active during G1 or S phases and whose dysregulation could impair cytokinesis. The small GTPase Rac1 is a negative regulator of cytokinesis that inhibits function of the contractile ring (Basant and Glotzer, 2018). Rac1 is active throughout the cell cycle except during a small window in mitosis, with its nadir of activity during anaphase and early telophase (Yoshizaki et al., 2003). Loss of Rac1 activity promotes cytokinesis (Canman et al., 2008), and overactivation of Rac1 impairs RhoA localization to the equatorial cell cortex (Bastos et al., 2012) and causes cytokinesis failure leading to formation of binucleate cells (Yoshizaki et al., 2004). To investigate whether BRAF^V600E^ affects Rac1 activity, we performed ELISA assays to detect active, GTP-bound Rac1 at various points following *BRAF^V600E^* expression and release from synchronization. As compared to control cells, GTP-bound Rac1 levels in *BRAF^V600E^*-expressing cells were higher upon release from synchronization and thereafter (Fig. 4B). To determine if higher Rac1 activity contributes to the *BRAF^V600E^*-induced formation of binucleate cells, we treated cells with the Rac1 inhibitors NSC2366 and EHT1864 and measured G1 tetraploid formation in *BRAF^V600E^*-expressing RPE-1 FUCCI cells. Treatment with either inhibitor suppressed the formation of tetraploid cells (Fig. 4C). Furthermore, treatment with either inhibitor led to the reestablishment of equatorial RhoA localization (Fig. 4D-E). Together these data indicate that BRAF^V600E^ acts through Rac1 to inhibit RhoA and cause failure of cytokinesis.

### Supernumerary centrosomes are observed in *BRAF^V600E^*-expressing cells

In assessing how Rac1 activity could be upregulated in *BRAF^V600E^*-expressing cells, we discovered that mitotic spindles in *BRAF^V600E^*-expressing cells showed structural organizations consistent with the presence of multiple, clustered centrosomes at the same spindle pole. Extra centrosomes increase microtubule nucleation, which in turn stimulates the activity of Rac1 (Godinho et al., 2014; Waterman-Storer et al., 1999). To determine whether extra centrosomes were present in *BRAF^V600E^*-expressing cells, we performed quantification of centrosomes via gamma tubulin staining in S phase cells. Here, we observed a significant increase in cells with more than 2 centrosomes in the *BRAF^V600E^*-induced population (Supplementary Fig. 4A, B). Gamma tubulin can stain fragments, derived from previously intact centrosomes, that can continue to nucleate microtubules (Rusan and Wadsworth, 2005). To address this possible artefact and confirm the presence of supernumerary centrosomal components in *BRAF^V600E^*-expressing cells, we stained for the centriolar marker Centrin-2 in mitotic cells (Fig. 5A). Supernumerary Centrin-2-positive centrioles (>4 centrioles per cell) were observed in 22% of *BRAF^V600E^*-expressing cells, which is an eight-fold increase as compared to control cells (Fig. 5B). In some cases, unpaired, single centrioles, indicative of a partially duplicated centrosome were observed (Supplementary Fig. 4C). Supernumerary centrioles were also present in *BRAF^V600E^*-expressing zebrafish melanocytes (Fig. 5C, D, Supplementary Fig. 4D, E). To determine whether supernumerary centrioles arose due to BRAF^V600E^-dependent MAPK signaling activity, we treated *BRAF^V600E^*-expressing cells with the MEK inhibitor trametinib and ERK inhibitor SCH772984. Both inhibitors suppressed the increase in centrioles (Fig. 5A, B), indicating that BRAF^V600E^-dependent overactivation of MAPK signaling led to supernumerary centrioles.

**Figure 5:**
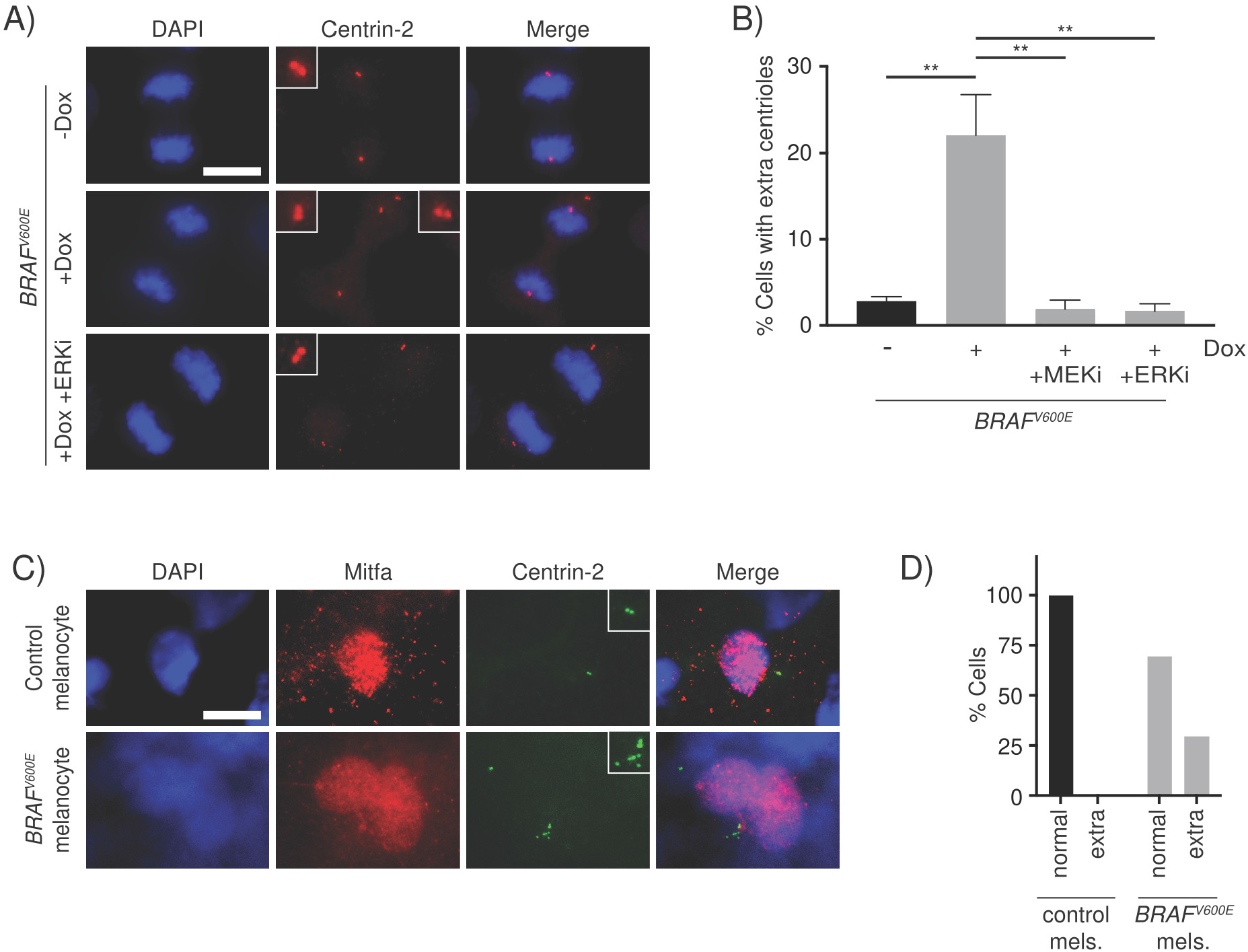
Supernumerary centrosomes are observed in *BRAF^V600E^*-expressing cells. A) DAPI and anti-CENTRIN-2 staining in control (-Dox) and *BRAF^V600E^*-expressing (+Dox) anaphase cells. Insets show centrioles at one pole. Images are maximum intensity projections of z-stacks. Scale bar = 7.5μM. B) Quantification of cells in mitosis with supernumerary (>4) centrioles. -Dox (n=131); +Dox (n=169); +Dox MEKi (n=85) and +Dox ERKi (n=91). Drugs were added coincident with Dox administration. Percent cells from 3 independent experiments is shown; one-way ANOVA with Tukey’s multiple comparisons test, ** p < 0.05. Error bars represent mean ± SEM. C) DAPI, anti-Mitfa and anti-CENTRIN-2 staining of control *Tg(mitfa:EGFP); alb(lf)* and *Tg(mitfa:EGFP); Tg(mitfa:BRAF^V600E^); alb(lf)* non-cycling zebrafish melanocytes. Scale bar = 7.5μM. Insets show centrioles. D) Percent cells with normal (2) and extra (>2) centrioles in control *Tg(mitfa:EGFP); alb(lf)* and *Tg(mitfa:EGFP); Tg(mitfa:BRAF^V600E^); alb(lf)* zebrafish melanocytes; chi squared test p = 0.000843.

### Supernumerary centrioles contribute to RhoA downregulation and *BRAF^V600E^*-induced WGD

Our findings indicated that BRAF^V600E^ caused RhoA downregulation and failure of cytokinesis as well as an increase in centrioles. We sought to determine whether the increase in centrioles contributed to RhoA downregulation and tetraploid cell formation. First, we examined if *BRAF^V600E^*-expressing cells with extra centrioles had reduced RhoA localization as compared to *BRAF^V600E^*-expressing cells with normal centrioles. *BRAF^V600E^*-expressing cells with a normal number and arrangement of centrioles had reduced RhoA staining; however, *BRAF^V600E^*-expressing cells with extra centrioles had an even greater reduction in RhoA localization (Fig. 6A, B). To establish that the reduced localization of RhoA in these cells was dependent on extra centrioles, we treated cells with centrinone, a PLK4 inhibitor that blocks centrosomal duplication (Wong et al., 2015). Centrinone treatment suppressed the formation of extra centrosomes and centrioles in *BRAF^V600E^*-expressing cells (Fig. 6C). The centrinone-treated *BRAF^V600E^*-expressing cells had higher RhoA equatorial localization as compared to control *BRAF^V600E^*-expressing cells (Fig. 6D, E). This recovery of RhoA localization was evident not only when centrinone treatment was coincident with *BRAF^V600E^* expression, but also when centrinone treatment was limited to late G1 and early S. Centrinone treatment also partially suppressed the *BRAF^V600E^*-driven formation of tetraploid cells (Fig. 6F). Thus, suppression of supernumerary centrioles in *BRAF^V600E^*-expressing cells recovered not only RhoA localization but also promoted cytokinesis. Taken together, these data indicate that the reduction of RhoA localization and cytokinesis in *BRAF^V600E^*-expressing cells is partially dependent on the *BRAF^V600E^*-driven increase in centrioles. Because the suppression of supernumerary centrioles in *BRAF^V600E^*-expressing cells did not fully restore RhoA localization and cytokinesis, a separate, centriole-independent mechanism driven by BRAF^V600E^ also reduces RhoA localization and cytokinesis.

**Figure 6:**
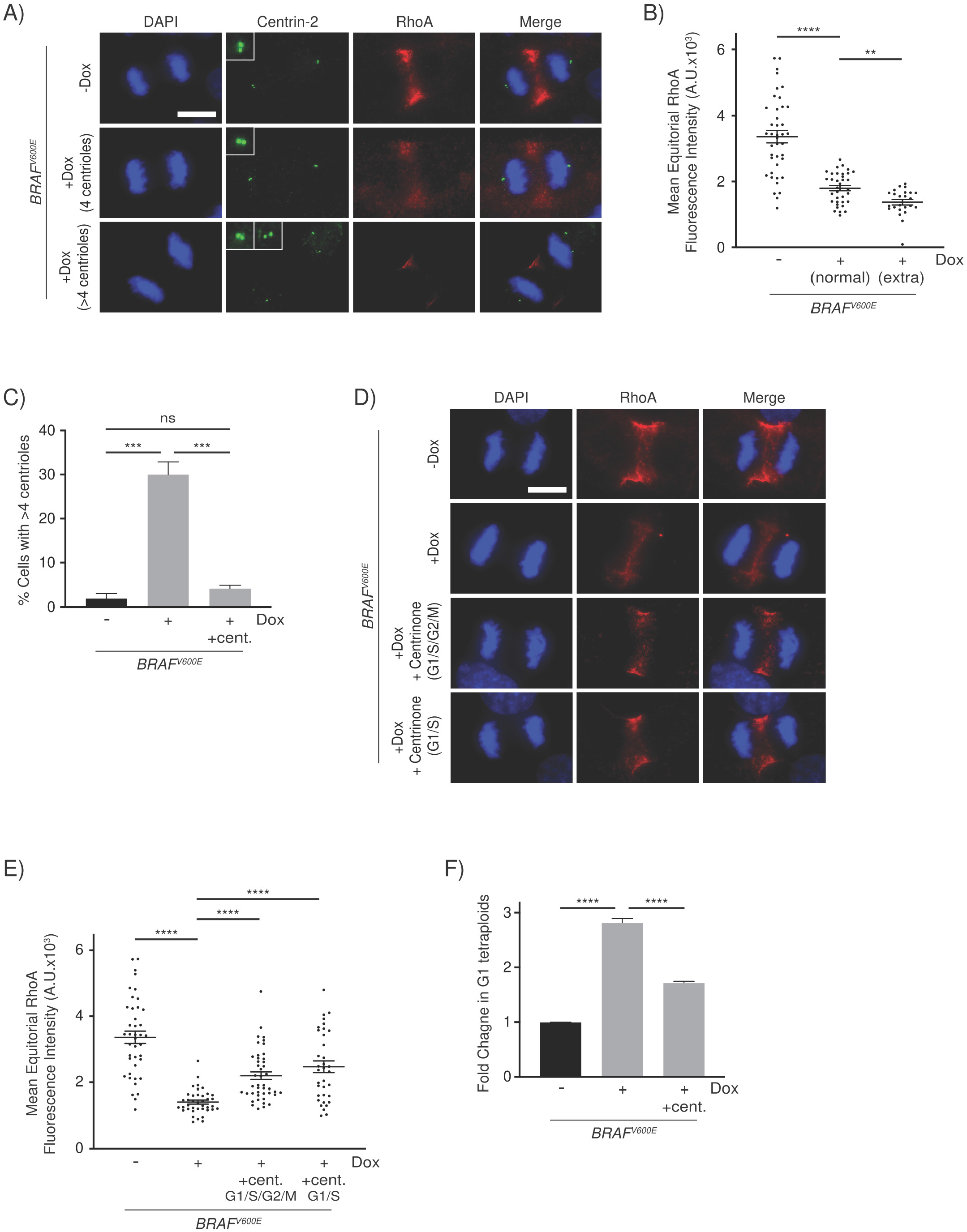
Supernumerary centrosomes contribute to RhoA downregulation and BRAF^V600E^-induced tetraploidy. A) DAPI, anti-CENTRIN-2 and anti-RhoA staining in control (-Dox) and *BRAF^V600E^*-expressing (+Dox) anaphase cells. Insets show centrioles at one pole. Images are maximum intensity projections of z-stacks. Scale bar = 7.5μM. B) Mean RhoA fluorescence intensity at the equator of control (n=30) and *BRAF^V600E^*-expressing anaphase cells with normal (n=54) and supernumerary (n=38) centrosomes. Fluorescence intensities (mean gray values) of the equator were measured by sum intensity projections of z-stacks. One-way ANOVA with Tukey’s multiple comparisons test, ** p < 0.01, **** p < 0.0001. Error bars represent mean ± SEM. C) Quantification of cells in mitosis with supernumerary (>4) centrioles. -Dox (n=119); +Dox (n=124) and +Dox +Centrinone (n=122). Centrinone was added coincident with Dox administration. Percent cells from 3 independent experiments is shown; one-way ANOVA with Tukey’s multiple comparisons test, *** p < 0.001, ns = not significant. Error bars represent mean ± SEM. D) DAPI and anti-RhoA staining in -*BRAF^V600E^* (-Dox) cells, *BRAF^V600E^*-expressing (+Dox) cells, and *BRAF^V600E^*-expressing (+Dox) cells treated with Centrinone. Centrinone was added coincident with Dox administration (G1/S/G2/M) or only during G1/S. Images are maximum intensity projections of z-stacks (0.20μM). Scale bar = 7.5μM. E) Mean RhoA fluorescence intensity at the equator of -*BRAF^V600E^* (-Dox) (n=32), *BRAF^V600E^*-expressing (+Dox) cells (n=40), and *BRAF^V600E^*-expressing (+Dox) cells treated with Centrinone coincident with Dox administration (G1/S/G2/M; n=44) or only during G1/S (n=41). Fluorescence intensities (mean gray values) of the equator were measured by sum intensity projections of z-stacks. One-way ANOVA with Tukey’s multiple comparisons test, **** p < 0.0001. Error bars represent mean ± SEM. F) Fold change in G1 tetraploids in control cells (-Dox), *BRAF^V600E^*-expressing cells (+Dox) and *BRAF^V600E^*-expressing cells treated with Centrinone (+Dox +cent.). Fold changes are expressed relative to the control cells. Fold change from 3 independent experiments is shown; One-way ANOVA with Tukey’s multiple comparisons test, **** p < 0.0001. Error bars represent mean ± SEM.

### p53 blocks cell cycle progression of *BRAF^V600E^*-induced tetraploid cells

Our finding that BRAF^V600E^ can cause failure of cytokinesis and lead to tetraploidy suggests that this may underlie the genome doubling events that are evident in BRAF^V600E^-mutated melanomas. However, for *BRAF^V600E^*-expressing cells to contribute to tumor formation, they would likely have to overcome the G1 phase arrest that tetraploid cells experience (Andreassen et al., 2001; Fujiwara et al., 2005). This tetraploid arrest can be triggered by Hippo pathway activation, which itself is activated by supernumerary centrosome-dependent activation of Rac1 (Ganem et al., 2014). Hippo pathway activation, in turn, activates p53, which has been shown to mediate arrest of tetraploid cells (Andreassen et al., 2001; Kuffer et al., 2013). Additionally, inactivation of p53 is strongly correlated with WGD in clinical samples, supporting the possibility of a p53-dependent arrest in tumors (Bielski et al., 2018).

To determine if *BRAF^V600E^*-expressing tetraploid cells were arrested, we first examined *Tg(mitfa:BRAF^V600E^)* as compared to *Tg(mitfa:BRAF^V600E^); p53(lf)* zebrafish melanocytes. Strikingly, the nuclei of *Tg(mitfa:BRAF^V600E^); p53(lf)* melanocytes were much larger than those of *Tg(mitfa:BRAF^V600E^)* melanocytes (Fig. 7A). Since DNA content is correlated with nuclear size (Heijo et al., 2020; Sher et al., 2013), we determined if *Tg(mitfa:BRAF^V600E^); p53(lf)* melanocyte nuclei had a higher DNA content than those of *Tg(mitfa:BRAF^V600E^)* melanocytes. Using DNA densitometry we found that, whereas *Tg(mitfa:BRAF^V600E^)* and *p53(lf)* nuclei were predominantly 2N, *Tg(mitfa:BRAF^V600E^); p53(lf)* nuclei were mostly 4N (Fig. 7B). This suggests that *BRAF^V600E^* expression causes an arrest of tetraploid cells that is dependent on p53. We also analyzed DNA content in zebrafish melanocytes by flow cytometry. GFP-positive melanocytes from *Tg(mitfa:EGFP); p53(lf); alb(lf)* animals were largely diploid as compared to melanocytes from *Tg(mitfa:EGFP); Tg(mitfa:BRAF^V600E^); p53(lf); alb(lf)* animals, most of which had a higher than 4N DNA content (Supplementary Fig. 5A), consistent with results from the nuclear DNA content densitometry analysis. These data suggest that, at least one and sometimes additional rounds of DNA replication can occur in the same melanocyte. The ploidy defects are dependent on both *BRAF^V600E^* expression and *p53* deficiency, mirroring the genetic synergism displayed by *Tg(mitfa:BRAF^V600E^)* and *p53(lf)* in melanoma formation.

**Figure 7:**
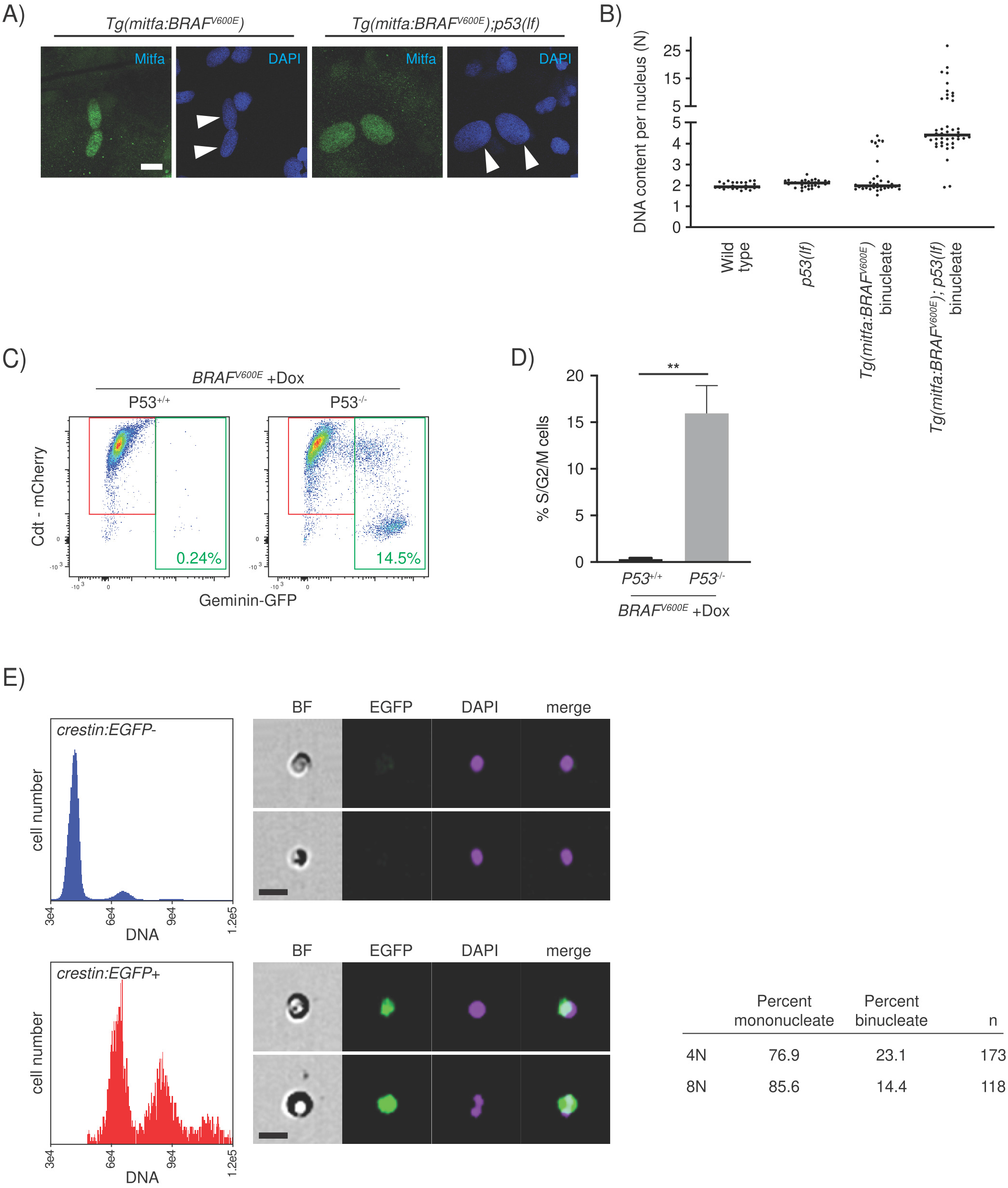
Progression of p53-mutant BRAF^V600E^-induced tetraploid cells. A) Anti-Mitfa and DAPI staining of nuclei in *Tg(mitfa:BRAF^V600E^)* and *Tg(mitfa:BRAF^V600E^); p53(lf)* melanocytes. Melanin pigment was bleached to more clearly visualize nuclear size. White arrowheads indicate 2 nuclei within a single melanocyte. Scale bar = 10μm. B) DNA content per nucleus as measured by confocal densitometry. Each data point represents a single nucleus. C) Flow cytometry plots of *BRAF^V600E^*-expressing (+Dox) p53 wild-type and p53 mutant tetraploid cells. Prior to analysis, G1 tetraploid cells were isolated and plated for 24 hours in the presence of nocodazole. Percentages of Geminin-GFP-positive cells in S/G2/M are indicated. D) Quantification of S/G2/M cells from flow cytometry analysis in (C). Percent cells from 3 independent experiments is shown; unpaired Student’s *t* test, ** p < 0.01. Error bars represent mean ± SEM. E) Flow cytometry and DNA content analysis of normal tissue (left, top) and nascent melanomas (left, bottom) from *Tg(crestin:EGFP); Tg(mitfa:BRAF^V600E^); p53(lf)* zebrafish. Brightfield, EGFP and DAPI images of single melanocytes are shown. Quantification of percent mononucleate and binucleate cells in the 4N and 8N peaks of nascent melanomas (right).

To more directly address a p53-dependent arrest, we used Crispr/Cas9-mediated genome editing to knock out P53 in our *BRAF^V600E^*-inducible RPE1-FUCCI cell clone. We isolated a clone in which both *P53* alleles were targeted and P53 protein expression abrogated (Supplementary Fig. 5B, C). These cells had DNA content profiles similar to P53 wild-type parental cells (Supplementary Fig. 5D, E). As compared to parental *BRAF^V600E^*-inducible RPE1-FUCCI cells, *BRAF^V600E^*-induclible *P53*^-/-^ RPE-1 FUCCI cells had a lower fraction of G1 tetraploids following *BRAF^V600E^* expression (Supplementary Fig. 5F, G), consistent with the notion that an arrest of G1 tetraploids was bypassed in *P53*^-/-^ cells. To determine if a P53-dependent arrest prevents *BRAF^V600E^*-expressing tetraploids from entering the cell cycle, we isolated G1 tetraploids after release from synchronization, cultured them for 24 hours, then assessed cell cycle progression. As measured by Geminin-GFP expression, *BRAF^V600E^*-expressing RPE1-FUCCI cells showed little to no progression (Fig. 7C, D). By contrast, nearly 15% of *BRAF^V600E^*-expressing p53^-/-^ RPE-1 FUCCI cells were Geminin-GFP-positive. Together these data indicate that *BRAF^V600E^*-induced tetraploid cells arrest in G1, and this arrest is alleviated by loss of P53.

### Nascent tumor cells in *BRAF^V600E^*-driven zebrafish melanomas are tetraploid and have higher ploidy

Our data indicate that BRAF^V600E^ can induce tetraploidy in melanocytes, and these melanocytes are prevented from progressing further by a p53-dependent block. Since *BRAF^V600E^* and loss of *p53* cooperate to form melanomas, and because *BRAF^V600E^* causes tetraploidy as seen in our *in vivo* and *in vitro* models, we were interested in understanding whether *BRAF^V600E^*-generated tetraploid cells serve as intermediates in melanomagenesis in our zebrafish model, potentially as cells of origin. This would be consistent with findings in tumor types with frequent WGD, in which tumors that have undergone a WGD event are thought to have undergone that event early in tumor progression (Jolly and Van Loo, 2018; Zack et al., 2013). To determine if tetraploid cells are present in the earliest stages of tumor formation, we took advantage of the zebrafish model in which early melanomas can be identified because of their reactivation of the neural crest gene *crestin* (Kaufman et al., 2016). Using a *Tg(mitfa:BRAF^V600E^); p53(lf); Tg(crestin:EGFP)* strain that marks tumor initiating cells, we identified zebrafish with early tumors (<20 cells), dissected these tumors and performed flow cytometry to determine DNA content of *crestin:EGFP*-positive tumor cells. The *crestin:EGFP*-positive cells were mostly tetraploid, and several octoploid cells were also observed (Fig. 7E). Notably, none of the cells were diploid, indicating that cells were tetraploid very early in tumor formation and it is likely that the cell of origin was tetraploid. Furthermore, the octoploid nascent tumor cells were mostly mononucleate (Fig. 7E) and thus different from the octoploid, binucleate melanocytes present in *Tg(mitfa:BRAF^V600E^); p53(lf)* strains. We speculate that these mononuclear, octoploid tumor cells arose from tetraploid cells of origin, and were cycling tetraploid cells in G2 and M phases. These data suggest that WGD can be present at the time of tumor initiation and potentially support tumor progression.

## Discussion

Our results indicate that BRAF^V600E^, and RAS/MAPK signaling in general, can cause the WGD that is a hallmark of melanomas and other tumor types. BRAF^V600E^ causes WGD by suppressing the activity of RhoA, leading to failure of cytokinesis. This suppression stems, in part, from supernumerary centrosomes that are formed as a result of BRAF^V600E^ activity. Supernumerary centrosomes are known to activate Rac1, and we found that Rac1 activity was necessary for BRAF^V600E^-dependent WGD. Taken together, our data support a model in which BRAF^V600E^ causes the formation of supernumerary centrosomes, leading to the activation of Rac1, which in turn causes inhibition of RhoA and failure of cytokinesis. The binucleate, tetraploid cells that result arrest in G0/G1 unless cells have an underlying mutation, in our models it is loss of P53, that abrogates the arrest and enables further cell cycle progression. Thus, in addition to stimulating cell cycle progression, suppressing cell death and providing other tumor-promoting activities, BRAF^V600E^ can cause WGD, which has been shown to support tumorigenesis and tumor progression.

A key component of this model is the formation of supernumerary centrosomes in *BRAF^V600E^*-expressing cells. The requirement for BRAF^V600E^ activity in G1/S coincides with the timing of centrosomal duplication in most cell types. Furthermore, gamma tubulin staining in *BRAF^V600E^*-expressing cells showed an increase in centrosome numbers in cells that had only progressed into S and G2 phases after *BRAF^V600E^* induction in G1. Together these observations suggest that the defect caused by BRAF^V600E^ is that of centrosomal overduplication. A similar phenotype has been observed upon *BRAF^V600E^* overexpression in established melanoma cell lines, although such cells had a high background of underlying centrosomal abnormalities (Cui et al., 2010; Liu et al., 2013). More recently, direct staining of melanoma samples suggests that supernumerary centrosomes in patient samples arise predominantly through overduplication (Denu et al., 2018). The mechanism by which BRAF^V600E^ could cause overduplication is not clear, although the ability of centrinone to suppress *BRAF^V600E^*-induced supernumerary centrosomes indicates that this mechanism is PLK4-dependent. As noted, the effect of supernumerary centrosomes only accounts for some of the RhoA reduction and cytokinesis failure upon *BRAF^V600E^* induction. Nonetheless, our findings together with the association of BRAF^V600E^ with centrosomal amplification in papillary thyroid, colorectal and other cancers suggests there may be a broad link between BRAF^V600E^-induced supernumerary centrosomes and WGD (Maric et al., 2011; Zhang et al., 2019).

Our findings begin to address the fate of cells that undergo WGD through cytokinesis failure. Zebrafish melanocytes and RPE-1 cells that have undergone cytokinesis failure arrest in G1 as binucleate tetraploid cells. However, in established zebrafish melanomas and late-stage human tumors that have previously experienced a WGD event, tumor cells go through the cell cycle, are predominantly mononuclear and do not indefinitely undergo cytokinesis failure. This raises the questions of how cells having just undergone a WGD event overcome the G1 block and, once they do, divide productively without creating giant, multinucleated cells. Newly-generated G1 binucleate tetraploid cells undergo a P53-dependent arrest, as evidenced by the progression into S phase of P53-mutant RPE-1 G1 tetraploids. The polyploid nature of *Tg(mitfa:BRAF^V600E^); p53(lf)* melanocytes indicates that this arrest also occurs in the zebrafish model. However, these zebrafish melanocytes are found as binucleates with most nuclei appearing to have gone through a single S phase without any further mitosis. This suggests that additional blocks may exist to prevent such cells from progressing further into the cell cycle, and such blocks would have to be overcome during tumorigenesis. If such cells were to progress and continue cycling, then conditions would have to exist to prevent them from stalling tumor growth as multinucleated cells. One possible condition is that the penetrance of BRAF^V600E^-induced cytokinesis failure is incomplete as in RPE-1 cells. Enough cytokinesis failure to enable WGD would be present, but not enough to prevent the amount of cell divisions required for tumor growth. Another possible condition is the attenuation of cytokinesis failure following a WGD event. RhoA activity has been shown to be upregulated in tumors (Fritz et al., 1999; Jung et al., 2020; Kakiuchi et al., 2014), and this could be an adaptation that prevents pervasive cytokinesis failure that would be detrimental to tumor progression.

Our finding that nascent zebrafish melanoma cells were tetraploid or had higher ploidy supports the notion that BRAF^V600E^-induced WGD can be present at the time of tumor initiation. In cancer types that have a high prevalence of WGD, such as melanoma, WGD in tumors frequently occurred early in tumor formation and might have been present at the time of tumor initiation and clonal outgrowth (Jolly and Van Loo, 2018; Zack et al., 2013). This raises the question of whether BRAF^V600E^, which is present in ~80% of benign nevi (Piris et al., 2015; Pollock et al., 2003), can cause WGD at the earliest stages of tumor formation and possibly in benign lesions. Although cells in common cutaneous nevi are primarily diploid (Newton et al., 1988; Winokur et al., 1990), several observations suggest that cytokinesis failure and associated WGD can occur in these cells. Cultures of nevocytes have been shown to contain binucleated cells (Gilchrest et al., 1986). Furthermore, the presence of binucleate and multinucleated giant cells in nevi is not uncommon (McKee, 2005), and polyploid cells are observed at a low fraction in many nevus samples (Skowronek et al., 1997). Together these observations suggest that some level of cytokinesis failure occurs in nevi and may be caused by BRAF^V600E^. The prevalence of early WGD in melanomas as compared to its relative absence in nevi, at least its absence when nevi begin clonal outgrowth, suggests there is some advantage to WGD in tumor formation. The ability to sample various pro-tumorigenic genomic configurations may underlie the benefit of WGD (Watkins et al., 2020) as would the ability of WGD to make haploid regions of a nascent tumor cell diploid and thus protect these regions from mutations incurred in essential genes (Lopez et al., 2020). Lastly, although the focus has been on WGD supporting tumor initiation, it is clear that many tumors analyzed bioinformatically, including the majority (~60%) of melanomas, show no evidence of having undergone WGD. However, this does not mean WGD is irrelevant in such tumors. Recent evidence suggests that WGD in melanomas can be late truncal and even occur privately in metastases (Birkeland et al., 2018; Vergara et al., 2021). Therefore, while WGD may not be necessary in all tumors for initiation, it nonetheless could play a role in disease progression. It is worth noting that the WGD observed in nevocytes, primary tumors and metastases overlaps with the presence of BRAF^V600E^ or other RAS/MAPK-activating mutations that could support WGD in such lesions.

## Acknowledgments

We thank Neil J. Ganem for sharing the RPE-1 FUCCI cell line, antibodies, and assistance with the live-cell imaging experiments; Kristyna Kotynkova for advice on RhoA and Anillin immunofluorescence staining; Paul Kaufman and Eric Campeau for the pLenti-CMV/TO neo/puro plasmids; Daryl Bosco for use of the Leica-DM inverted microscope; Desiree Baron for imaging and microscopy advice; Charles Kaufman for the *Tg(crestin:EGFP)* zebrafish strain; Melissa Guerin for help with fish genotyping; Patrick White, Ed Jaskolski and the staff at the UMMS Animal Medicine Department for fish care; Tammy Krumpoch and Susanne Pechhold at the UMMS Flow Cytometry Core for guidance and assistance in performing flow cytometry and FACS experiments. This research was supported by NIAMS AR063850 to C.J.C.

## Author Contributions

R.D. and C.J.C. conceptualized the study, designed the *in vitro* experiments, and wrote the manuscript. R.D. and C.J.C. designed all the zebrafish experiments. R.D. performed all the *in vitro* biological assays, immunofluorescence staining, flow cytometry and image analysis. M.A.V. and N.J.G. performed the live-cell imaging experiments and R.D. analyzed the data. N.J.G. provided critical reagents.

## Methods

### Zebrafish strains and husbandry

Strains were maintained at 28°C as described by Westerfield (Westerfield, 2000). AB was used as the wild-type strain. The following mutations were used: *p53(zdf1)* (Berghmans et al., 2005), *mitfa(w2) (Lister et al., 1999), alb(b4) (Chakrabarti et al., 1983)* and are designated throughout as “*lf*’ for loss-of-function mutation. The integrated transgene *Tg(mitfa:BRAF^V600E^)* expresses oncogenic human *BRAF^V600E^* under the control of the zebrafish pigment cell-specific *mitfa* promoter (Patton et al., 2005).

### Cell line generation

RPE-1 FUCCI cells were a gift from Neil Ganem (Boston University). RPE-1 cells were grown in DMEM: F12 media containing 10% tetracycline-free FBS, 100 IU/ml penicillin, and 100 μg/ml streptomycin. Cells were maintained at 37°C with 5% CO_2_ atmosphere. TetR was expressed in cells using pLenti CMV TetR Blast (Addgene #17492). Transduced cells were selected with 5μg/ml blasticidin and bulk cells expressing the TetR protein were clonally isolated in 96-well plates using FACS (BD-Aria). *BRAF^V600E^* and *BRAF^WT^* were expressed in cells using Gateway-compatible lentiviral constructs (pLenti CMV/TO puro DEST, pLenti CMV/TO neo DEST; gifts from Eric Campeau and Paul Kaufman (Campeau et al., 2009)). Lentivirus carrying *BRAF^WT^* or *BRAF^V600E^* construct was generated by transfection of HEK-293T cells, with appropriate packaging plasmids (pMD2.G and psPAX2) using lipofectamine 2000, according to the manufacturer’s instructions. RPE-1 cells were infected for 48 hrs with virus carrying a BRAF construct in the presence of 8 μg/ml polybrene, washed, and allowed to recover for 24hrs before selection. Cells were selected with the appropriate antibiotic selection marker (5μg/ml puromycin or 1mg/ml neomycin) and clonally isolated in 96-well plates using FACS on the BD-Aria. Doxycycline was used at 1μg/ml to induce the expression of *BRAF*.

### Synchronization and ploidy analysis

To turn on *BRAF^V600E^* in a synchronized cell population, we serum starved RPE-1 FUCCI cells with 0.1% serum for 48hrs. Serum-starved cells were released into media with 10% serum until they reached the next G1 (~33hrs). *BRAF^V600E^* was turned on using doxycycline, and cells were resynchronized at G1/S using thymidine (2.5mM). Following thymidine washout cells were allowed to progress through the cell cycle, and ploidy was analyzed in next G1 (16hrs after thymidine release) using Hoechst incorporation (2.5μg/ml) in hCdt1-mcherry expressing cells by flow cytometry (BD-LSR). Additional information is provided in Extended Experimental Procedures. For immunofluorescence experiments of mitotic cells, cells were fixed at 12hrs after thymidine release.

### Live cell imaging

For live cell imaging, RPE-1 H2B-GFP-transfected cells were grown on glass-bottom 12-well tissue culture dishes (MatTek) and imaged on a Nikon TE2000E inverted microscope equipped with a cooled Hamamatsu Orca-ER CCD 11 camera and the Nikon Perfect Focus system. The microscope was enclosed within an incubation chamber that maintained an atmosphere of 37°C and 3-5% humidified CO_2_. GFP and phase-contrast images were captured at multiple locations every 5 minutes with either 20X or 40X Nikon Plan Fluor objectives. All captured images were analyzed using NIS-Elements software. For imaging, cells were synchronized with thymidine and imaged for 48 hours after release. Each cell going through mitosis was observed and cytokinesis was scored by phase contrast imaging. To verify cytokinesis failure and to confirm that the nuclei stayed within one cell, cells were carefully traced for at least 10 frames. To quantify mitotic duration, each frame beginning from the start of nuclear envelope breakdown to anaphase onset was counted.

### Immunofluorescence and imaging

RPE-1 BRAF^V600E^ cells were grown directly on collagen IV (Sigma) -coated coverslips, fixed in 3.7% formalin, permeabilized using 0.1% Triton X-100, and treated with 0.1% SDS. They were blocked in 1% BSA and then incubated with primary antibody diluted in blocking solution in a humidity chamber at 4°C overnight, washed with 1X PBS, and incubated with secondary antibody. Cells were mounted using mounting media containing DAPI (Vector Laboratories). All secondary antibodies were Alexa Fluor conjugates (488, 555, and 647) (Thermofisher) used at a 1:500 dilution. All images were acquired on a Leica DM 600 inverted microscope at 100X magnification. Z stack images were taken at 0.20μM each. For melanocyte binucleate counts and immunofluorescence staining, zebrafish adult dorsal scales were fixed using 4% paraformaldehyde for two hours, washed with PBST (PBS+ 0.1% Triton X) and water then blocked in 1%BSA/PBS for 30 minutes prior to primary antibody incubation. Affinity-purified anti-Mitfa antibodies were used at a 1:100 dilution for staining. All images were acquired on a Nikon Eclipse Ti A1R-A1 confocal microscope. Additional information is provided in Extended Experimental Procedures.

### RhoA and Rac1 activation assays

RPE-FUCCI cells were synchronized as described above, and lysates were collected at 0, 4, 8, 12hrs post-release from thymidine. GTP-bound RhoA and Rac1 were immunoprecipitated from cells according to manufacturers’ protocols (BK-036, BK-128, Cytoskeleton).

### Western blotting

Cells were lysed in ice-cold RIPA buffer containing a Complete protease inhibitor tablet (Roche). Protein concentration was measured using the Pierce BCA Protein Assay Kit (Life Technologies). Samples were run on 10% polyacrylamide gels, transferred, and developed using fluorophore-conjugated antibodies (LI-COR). Antibodies against the following proteins were used: pMEK1/2 (S217/221), MEK1/2 and p44/42 MAPK (ERK1/2), alpha-Tubulin (Cell Signaling Technology 9154, 8727, 4695, 3873, respectively); pERK (Sigma m8159); BRAF^V600E^ (Spring Bioscience E19290); BRAF (Millipore 10146); IRDye 800CW Donkey anti-Rabbit and IRDye 680RD Goat anti-Mouse (LI-COR 926-32213 and 926-68070, respectively). Additional information is provided in Extended Experimental Procedures.

### Cyclin D1 staining for flow cytometry

RPE-1 FUCCI cells were trypsinized, washed with 1XPBS and fixed in 2% PFA at RT for 15min. Cells were then washed with PBS/1%FBS and permeabilized with 100% methanol (for cyclin D1). Cells were washed with PBS, incubated with Alexa Fluor 647-conjugated primary antibodies for 60 minutes at room temperature, washed and stained with 1:1000 DAPI for 15 minutes before flow cytometry (BD-LSR).

### CRISPR cell line generation

Oligos targeting p53 were annealed and inserted into lentiCRISPR v2 (gift of Feng Zhang) as described previously (Sanjana et al., 2014; Shalem et al., 2014) The lentiviral packaging plasmids pMD2.G (Addgene plasmid #12259) and psPAX2 (Addgene plasmid #12260) were used for transfection using lipofectamine. For lentiviral transduction, cells (300,000) were plated in a 6 well plate the day before transduction. Lentivirus was harvested and added to OptiMEM supplemented with 8 μg/mL polybrene. Media was changed 24 h after transduction to remove polybrene. Media supplemented with 5 μg/mL puromycin (Sigma Aldrich) or was changed 48h after transduction to select lentiCRISPRv2 transduced cells. Following transduction with p53-targeting lentiCRISPRv2 containing lentivirus, antibiotic-resistant cells were selected then clonally isolated by FACS in 96 well plates. Clones were then assayed for indels via the surveyor assay (706020,IDT). Cells that showed indels were then cloned into the pGEM-T Easy vector (Promega), and colonies were picked using blue/white screening then sequenced. The p53 gRNA sequence was CCCCGGACGATATTGAACAA. Primers used for sequencing indels were as follows:

Forward - 5’ GTAAGGACAAGGGTTGGGCT 3’

Reverse - 5’ GAAGTCTCATGGAAGCCAGC 3’

### Flow cytometry of zebrafish melanocytes, normal tissue and tumors

To isolate zebrafish melanocytes for flow cytometry, 4-to 6-month old fish were treated for 5 minutes with the anesthetic tricaine methanesulfonate. Scales were plucked from the dorsal anterior region of fish and put in 0.9X PBS. Cells were dissociated using TH liberase (Roche) for 30 mins by constant agitation at 37°C. Cells were immediately fixed for 2 hours in 4% paraformaldehyde. Following fixation, cells were washed and permeabilized 3X with 0.1% Triton X/PBS and immediately stained with DAPI (1:1000). Cells were then filtered through a 40μM mesh filter and spun down at 2000rpm. Analysis for GFP+ and DAPI+ cells was performed on the Amnis Flowsight.

For ploidy analysis of zebrafish tumors, melanomas from *Tg(mitfa:EGFP)(mitfa:BRAF^V600E^); p53(lf);alb* animals were removed, homogenized in PBS, and stained and analyzed in the same manner as described above. For normal tissue analysis, the caudal peduncle and fin were dissected and homogenized.

### Confocal densitometry

Zebrafish scales were obtained and stained with anti-Mitfa antibody and DAPI as described above. Melanin pigment interferes with quantitative UV-based imaging so scales were bleached prior to staining. Mitfa-positive melanocyte nuclei were identified, and Z stacks of the DAPI signal of these nuclei were obtained. The same distance between Z slices (0.37μM) and pixel intensity lookup table were used for each nucleus measured. In analyzing each slice, nuclear boundaries were specified, and pixel intensity values within the nuclear area measured. Pixel intensities for all slices of one nucleus were summed to derive a raw DNA content. Ploidy was estimated using nearby Mitfa-negative nuclei as 2N controls. Nuclei in the same binucleate cell are typically arranged as mirror-image pairs, and this orientation allowed us to identify nuclei of binucleate cells in the absence of melanin pigment.

### Melanocyte density assay

4- to 6-month old fish were treated for 5 minutes with the anesthetic tricaine methanesulfonate and epinephrine, which contracts melanosomes to the central cell body of melanocytes (Goodrich and Nichols, 1931), thereby resolving overlapping cells. Scales were plucked from the dorsal anterior region of fish from the scale rows adjacent to the dorsal midline row. Scales were immediately fixed for ≥30 minutes in 4% paraformaldehyde. After fixation, scales were flat mounted and melanocytes counted. Area was estimated by multiplying maximal antero-posterior and left-right distances of the scale.

### Statistical analysis

Significance calculations were performed on samples collected in a minimum of biological triplicate. *P* values from two-tailed Student’s *t* tests or ANOVA were calculated for all comparisons of continuous variables. All further significance tests were performed in GraphPad Prism (v8.0). A *P* value < 0.05 was considered significant.

## Extended Experimental Procedures

### Drug treatments

Cells were synchronized as described with serum starvation and thymidine arrest. Treatment of cells with small molecule compounds was performed as follows: G1/S/G2/M – drug was added 33 hours following serum addition and coincident with doxycycline; S/G2/M – drug was added following thymidine washout; G1/S – drug was added 39 hours following serum addition and washed off 12 hours later; G1 – drug was added 33 hours following serum addition and washed off 6 hours later; early G1 – drug was added XX hours following serum addition and washed off XX hours later. Small molecule inhibitors used were – Vemurafenib, 1μM (S1267, Selleck Chemicals), Trametinib 50nM (S2673, Selleck Chemicals), SCH772984 50nM (S1701, Selleck Chemicals), PLX7904 1μM (S7964, Selleck Chemicals), PLX8394 1μM (HY-18972, MCE), Centrinone 300nM (5687, Tocris), NSC2366 5μM (13196, Cayman Chem), EHT1864 5μM (17258, Cayman Chem), LPA 1 μM (BML-LP100-0005, Enzo), S1P 1μM (62570, Cayman Chem), Dihydrocytochalasin B 4μM (D1641, Sigma).

### Immunofluorescence and antibodies

For RhoA staining, cells were fixed in 10% Trichloroacetic acid (TCA) (T6399, Sigma) on ice for 15 min, washed with PBS/30nM glycine, then permeabilized with 0.2% Triton X/PBS/30nM glycine for 10 mins on a rocking platform. Cells were then washed with PBS/30nM glycine, blocked with 3% BSA/PBS/0.01% Triton X-100 for 1hr at room temperature before staining with primary antibody in a humidity chamber at 4°C overnight. For Anillin Staining, cells were fixed in ice cold methanol at −20°C for 2 hrs. Cells were then washed with PBST (PBS/0.01% Triton X-100), permeabilized with 0.2% Triton X-100 in PBS for 10 min, washed with PBST and then incubated with primary antibody in a humidity chamber at 4°C overnight. For Centrin-2 staining, cells were fixed in ice cold 100% methanol at −20°C, permeabilized with 0.2% TritonX/PBS and blocked in 1%BSA/PBS before incubating with primary antibody (1:100) in a humidity chamber at 4°C, overnight. Primary antibodies and dilutions used for immunofluorescence were: RhoA 1:75 (sc-418, SCBT), Anillin 1:100 (PA5-28645, Thermofisher), CENTRIN-2 1:100 (04-1624, Millipore), *γ* – TUBULIN 1:500 (T6557, Sigma), *α*-tubulin DM1A 1:2000 (3873S, Cell Signaling). Primary antibodies and dilutions used for western blot were: TetR, 1:1000 (631132, Takara), RhoA, 1:500 (ARH05, Cytoskeleton), RAC1 1:500 (ARC03, Cytoskeleton), GAPDH 1:5000 (AM4300, Thermofisher), p53 1:1000 (sc-126, SCBT), TUBULIN 1:1000(2144S, Cell Signaling), CYCLIN D1, 1:100 (NBP2-33138AF647, Novus Bio), Rabbit IgG Isotype control, 1:100 (3452S, Cell Signaling), HA tag, 1:100 (IC6875R, R&D systems), Mouse IgG1 Isotype control 1:100 (IC002R, R&D systems).

### RhoA and Anillin intensity measurements

Fluorescence intensities were quantified using ImageJ as previously described (Kotynkova et al., 2016). Briefly, Z-stack images were loaded on ImageJ and the sum intensity projection was obtained using the Z-stack function. RhoA and Anillin equatorial fluorescence intensities were obtained by measuring the intensity profile of the fluorescence signal along a line manually placed along the cell equator, parallel to the anaphase DNA position. The mean fluorescence intensity was obtained by averaging the two intensity values at each side of the furrow. The mean background signal was obtained by averaging the signal of three manually selected circular regions with a diameter of 50 pixels outside of the cell and the value was subtracted from the equatorial intensities.

### Growth and apoptosis assays

To perform growth curves, 20,000 cells were plated in 12-well plates and appropriate samples were treated with inhibitors and doxycycline. Cells were counted every 24hrs over 4 days using the hemocytometer. Cell death following inhibitor treatment was assessed by the caspase glow assay according to the manufacturer’s instructions (G8090,Promega).

## Notes

The authors declare no potential conflicts of interest.

### Competing Interest Statement

The authors have declared no competing interest.

