## Supplemental Data for "Oncogenic BRAF Induces Whole-Genome Doubling Through Suppression of Cytokinesis"

#### SUPPLEMENTARY FIGURE 1

A)

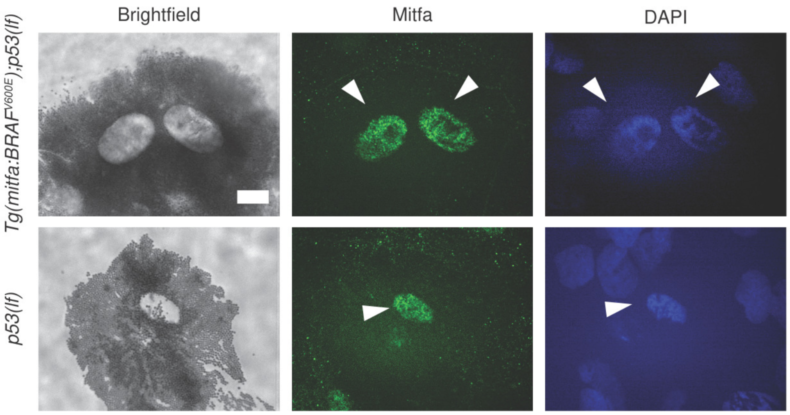


B)

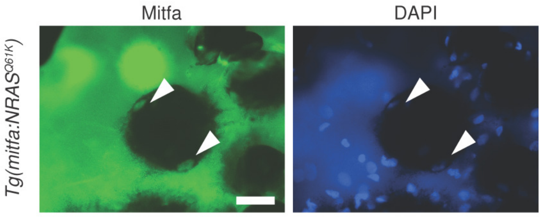

**Supplementary Figure 1**

A) Images from brightfield (left), anti-Mitfa (middle) and DAPI (right) staining of a single *Tg(mitfa:BRAF$^{V600E}$); p53(lf)* (top) or *p53(lf)* (bottom) epidermal melanocyte. Only the melanocyte nuclei stain positively for Mitfa. White arrowheads indicate nuclei within a single melanocyte. Scale bar = 5μm.

B) Images from anti-Mitfa (left) and DAPI (right) staining of a single *Tg(mitfa:NRASQ61K)* epidermal melanocyte. White arrowheads indicate nuclei within a single melanocyte. Scale bar = 20μm.

### SUPPLEMENTARY FIGURE 2

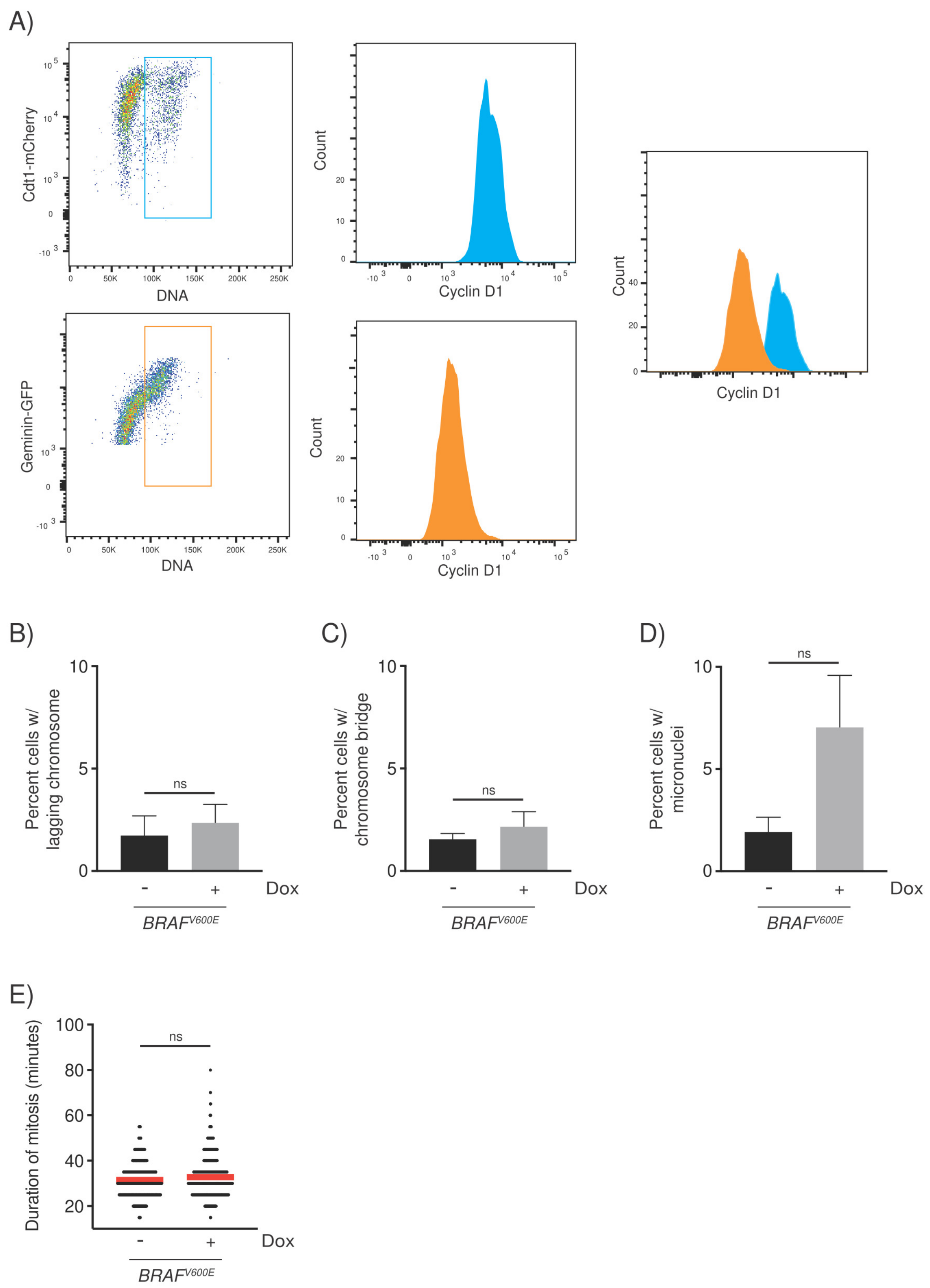

**Supplementary Figure 2**

A) Flow cytometry plots of Cyclin D1 analysis of RPE-1 FUCCI cells in G1 vs S/G2/M phases. Gating of Cdt1-mCherry-positive 4N cells (top left) and Geminin-GFP-positive 4N cells (bottom left). Cyclin D1 staining of 4N populations (middle), and an overlay of Cyclin D1 histograms (right) comparing Cdt1-mCherry-positive 4N cells (blue) and Geminin-GFP-positive 4N cells (orange).

B) Fold change in percent cells with lagging chromosomes. Fold change from 3 independent experiments is shown; unpaired Student's *t* test, ns = not significant. Error bars represent mean ± SEM.

C) Fold change in percent cells with chromosome bridges. Fold change from 3 independent experiments is shown; unpaired Student's *t* test, ns = not significant. Error bars represent mean ± SEM.

D) Fold change in percent cells with micronuclei. Fold change from 3 independent experiments is shown; unpaired Student's *t* test, ns = not significant. Error bars represent mean ± SEM.

E) Duration of mitosis in control (-Dox) and $BRAF^{V600E}$-expressing (+Dox) cells as quantified from live-cell imaging experiments. Each data point represents a single cell; unpaired Student's *t* test, ns = not significant. Mean duration is indicated.

#### SUPPLEMENTARY FIGURE 3

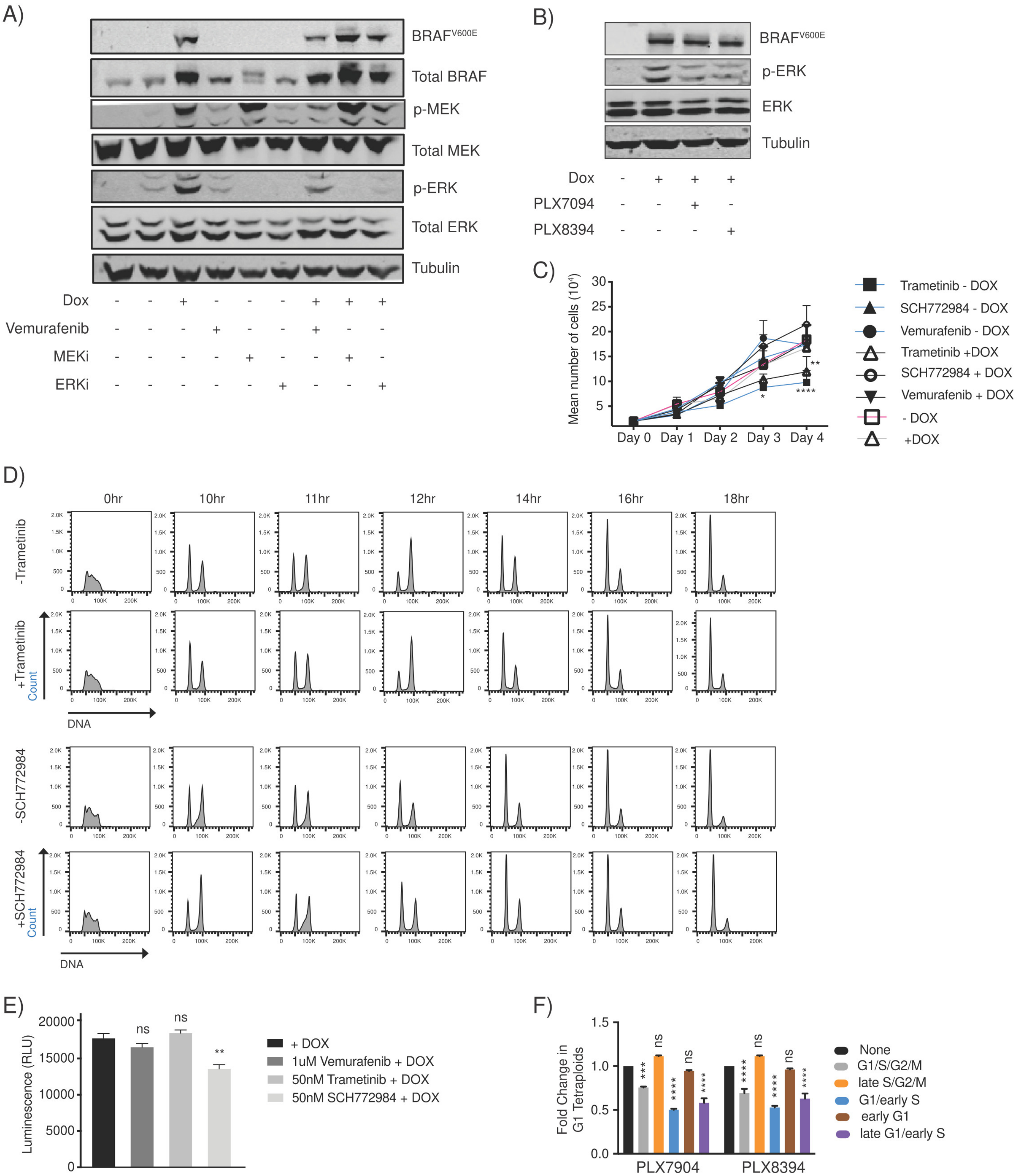

**Supplementary Figure 3**

A) Western blot of phosphorylated MEK and phosphorylated ERK upon treatment with BRAF, MEK and ERK inhibitors. Synchronized control (-Dox) or *BRAF*$^{V600E}$-expressing (+Dox) RPE-1 FUCCI cells were harvested 48hrs following incubation with inhibitors. Inhibitors used were the BRAF inhibitor Vemurafenib, MEK inhibitor Trametinib (MEKi) and ERK inhibitor SCH772984 (ERKi). Tubulin was used as a loading control.
B) Western blot of phosphorylated ERK upon treatment with BRAF paradox-breaking inhibitors. Synchronized control (-Dox) or *BRAF*$^{V600E}$-expressing (+Dox) RPE-1 FUCCI cells were harvested 48hrs following incubation with inhibitors. Inhibitors used were the BRAF paradox-breaking inhibitors PLX7094 and PLX8394. Tubulin was used as a loading control.
C) Flow cytometry plots showing cell cycle progression of control untreated cells and cells treated with Trametinib (MEKi) and SCH772984 (ERKi). The 0hr timepoint corresponds to when cells were released from thymidine block.
D) Proliferation curves of control untreated cells and cells treated with Trametinib (MEKi), SCH772984 (ERKi) and Vemurafenib (BRAFi). Cells were counted every 24hrs to record cell counts.
E) Caspase-glo assay to quantify apoptosis of control untreated cells and cells treated with Trametinib (MEKi), SCH772984 (ERKi) and Vemurafenib (BRAFi).
F) Fold change in G1 tetraploids following BRAF paradox-breaking inhibitor treatment relative to control (+*BRAF*$^{V600E}$ No Drug) samples. Inhibitors were added at indicated timepoints. Fold changes from 3 independent experiments are shown; unpaired Student's *t* test, * $p < 0.05$, ** $p < 0.01$, ***$p < 0.001$, ****$p < 0.0001$, ns = not significant. Error bars represent mean ± SEM.

#### SUPPLEMENTARY FIGURE 4

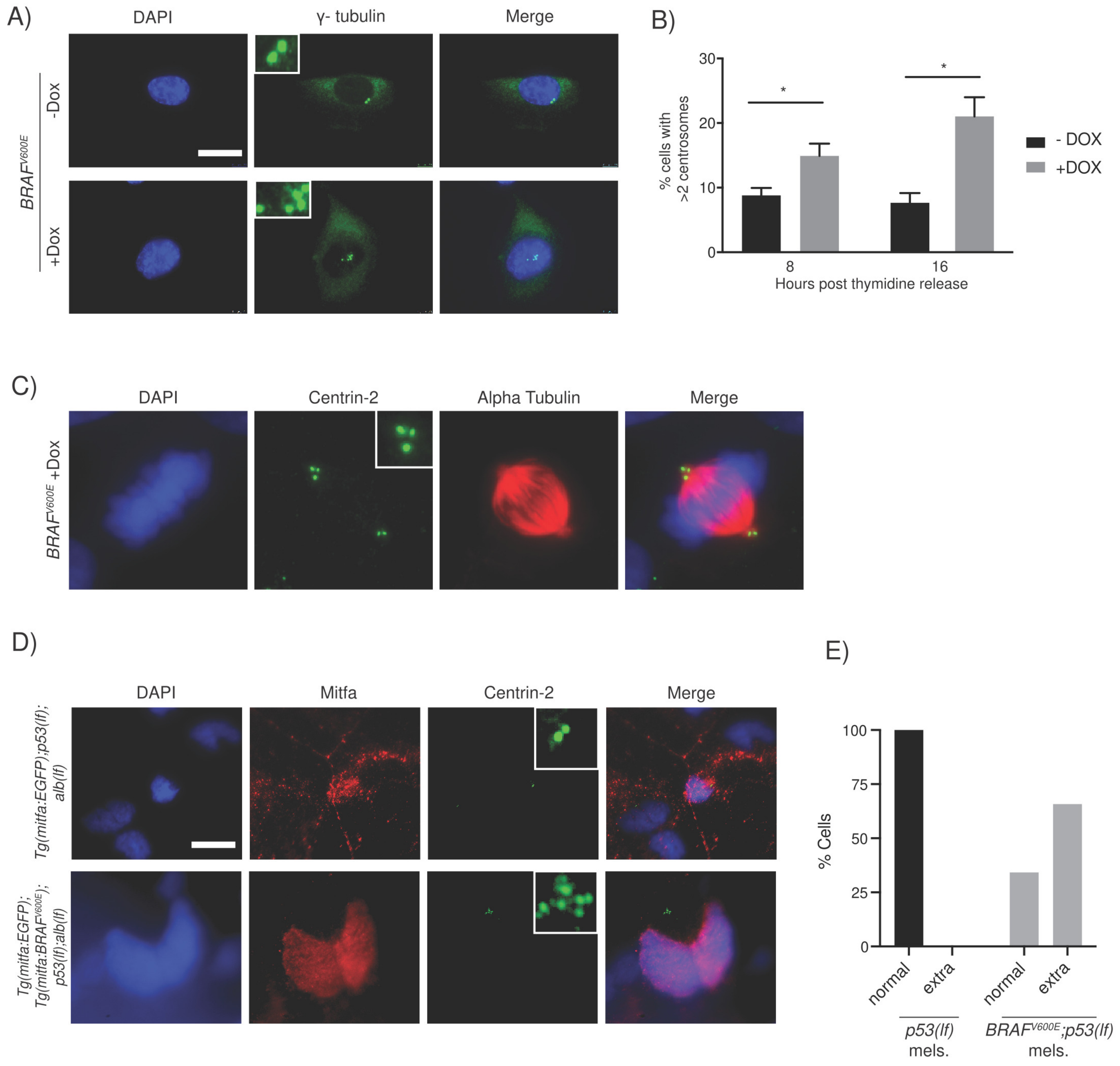

**Supplementary Figure 4**

A) DAPI and anti-gamma tubulin staining of control (-Dox) and *BRAF*$^{V600E}$-expressing (+Dox) cells. Images shown were taken of S/G2 cells at 8hrs post thymidine release. Insets show centrosomes. Scale bar= 7.5μM.

B) Cells with >2 centrosomes were quantified at 8hrs (n= 1105 for -Dox and n= 1093 for +Dox) and 16hrs (n= 1445 for -Dox and n=1262 for +Dox) post thymidine release. Percent cells from 3 independent experiments is shown; unpaired Student's *t* test, * $p < 0.05$. Error bars represent mean ± SEM.

C) DAPI and anti-centrin-2 staining of *BRAF*$^{V600E}$-expressing (+Dox) mitotic cells. Insets show centrioles at one pole. Merged image shows clustering of supernumerary centrosomes at one spindle pole. Scale bar = 7.5μM.

D) DAPI, Mitfa and Centrin-2 staining of *Tg(mitfa:EGFP); p53(lf); alb(lf)* and *Tg(mitfa:EGFP); Tg(mitfa:BRAF*$^{V600E}$*); p53(lf); alb(lf)* in non-cycling zebrafish melanocytes. Scale bar = 7.5μM. Insets show centrioles.

E) Percent cells with normal and extra centrioles in control *Tg(mitfa:EGFP); p53(lf); alb(lf)* and *Tg(mitfa:EGFP); Tg(mitfa:BRAF*$^{V600E}$*); p53(lf); alb(lf)* zebrafish melanocytes. Normal centrioles are 2 per nucleus, so extra centrioles are >2 in mononuclear *Tg(mitfa:EGFP); p53(lf); alb(lf)* cells and >4 in binucleate *Tg(mitfa:EGFP); Tg(mitfa:BRAF*$^{V600E}$*); p53(lf); alb(lf)* cells; chi squared test p = 0.000009

#### SUPPLEMENTARY FIGURE 5

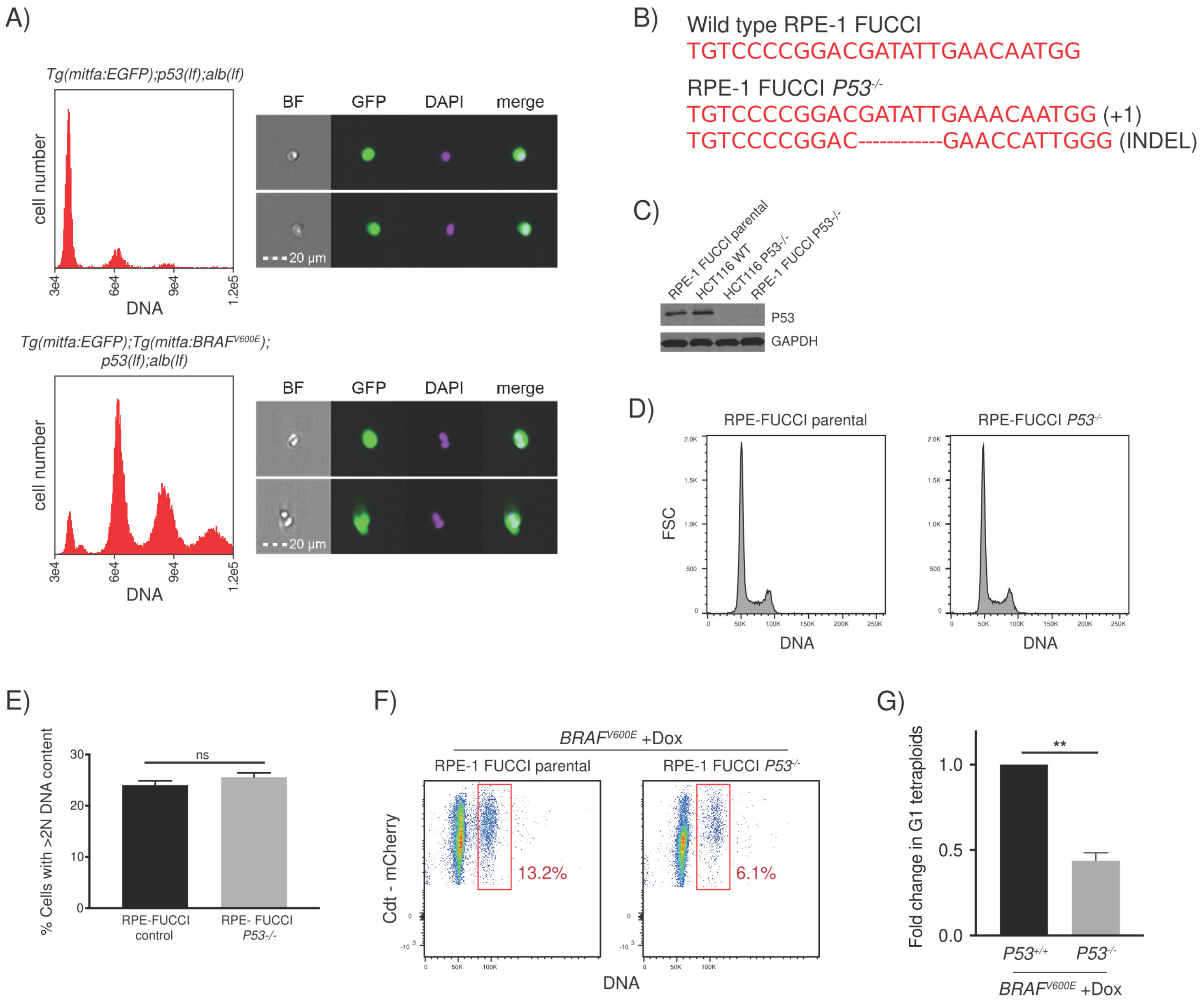

**Supplementary Figure 5**

A) Flow cytometry and DNA content analysis of *Tg(mitfa:EGFP); p53(lf); alb(lf)* and *Tg(mitfa:EGFP); Tg(mitfa:BRAF$^{V600E}$); p53(lf); alb(lf)* melanocytes with brightfield, EGFP and DAPI images of single melanocytes.

B) Genotype of Crispr/Cas9-targeted RPE-1 FUCCI P53$^{-/-}$ cells.

C) Western blot of P53 in RPE-1 FUCCI P53$^{+/+}$ compared to RPE-1 FUCCI P53$^{-/-}$ cells. HCT116 P53$^{+/+}$ and HCT116 P53$^{-/-}$ cells were used as controls. GAPDH was a loading control.

D) Flow cytometry histograms comparing DNA content of unsynchronized RPE-1 FUCCI control cells and RPE-1 FUCCI P53$^{-/-}$ cells.

E) Percent cells with >2N DNA content; unpaired Student's *t* test, ns = not significant. Error bars represent mean ± SEM.

F) Flow cytometry plots of *BRAF$^{V600E}$*-expressing (+Dox) RPE-1 FUCCI parental and RPE-1 FUCCI P53$^{-/-}$ cells. Tetraploid cells accumulating in G1 were quantified based on Cdt1-mCherry positivity and Hoechst incorporation. Percentages of G1 tetraploid cells in control and *BRAF$^{V600E}$*-expressing cultures are indicated.

G) Fold change in G1 tetraploids of *BRAF$^{V600E}$*-expressing (+Dox) RPE-1 FUCCI P53-/- cells relative to control parental cells. Fold change from 3 independent experiments is shown; unpaired Student's *t* test, ** $p < 0.01$. Error bars represent mean ± SEM.
